# Glycoprotein VI is Critical for the Detection and Progression of Abdominal Aortic Aneurysms

**DOI:** 10.1101/2023.07.02.547361

**Authors:** Tyler W. Benson, Mindy M. Pike, Anthony Spuzzillo, Sarah M. Hicks, Michael Pham, Doran S. Mix, Seth I. Brunner, Caris Wadding-Lee, Kelsey A. Conrad, Hannah M. Russell, Courtney Jennings, Taylor M. Coughlin, Anu Aggarwal, Sean Lyden, Kevin Mani, Martin Björck, Anders Wanhainen, Rohan Bhandari, Loren Lipworth-Elliot, Cassianne Robinson-Cohen, Francis J. Caputo, Sharon Shim, Todd L. Edwards, Michael Tranter, Elizabeth E. Gardiner, Nigel Mackman, Scott J. Cameron, A. Phillip Owens

## Abstract

A common feature in patients with abdominal aortic aneurysms (AAA) is the formation of a nonocclusive intraluminal thrombus (ILT) in regions of aortic dilation. Platelets are known to maintain hemostasis and propagate thrombosis through several redundant activation mechanisms, yet the role of platelet activation in the pathogenesis of AAA associated ILT is still poorly understood. Thus, we sought to investigate how platelet activation impacts the pathogenesis of AAA. Using RNA-sequencing, we identify that the platelet-associated transcripts are significantly enriched in the ILT compared to the adjacent aneurysm wall and healthy control aortas. We found that the platelet specific receptor glycoprotein VI (GPVI) is among the top enriched genes in AAA ILT and is increased on the platelet surface of AAA patients. Examination of a specific indicator of platelet activity, soluble GPVI (sGPVI), in two independent AAA patient cohorts is highly predictive of a AAA diagnosis and associates more strongly with aneurysm growth rate when compared to D-dimer in humans. Finally, intervention with the anti-GPVI antibody (J) in mice with established aneurysms blunted the progression of AAA in two independent mouse models. In conclusion, we show that levels of sGPVI in humans can predict a diagnosis of AAA and AAA growth rate, which may be critical in the identification of high-risk patients. We also identify GPVI as a novel platelet-specific AAA therapeutic target, with minimal risk of adverse bleeding complications, where none currently exist.

**KEY POINTS:**

- Soluble glycoprotein VI, which is a platelet-derived blood biomarker, predicts a diagnosis of AAA, with high sensitivity and specificity in distinguishing patients with fast from slow-growing AAA.
- Blockade of glycoprotein VI in mice with established aneurysms reduces AAA progression and mortality, indicating therapeutic potential.

## INTRODUCTION

Abdominal aortic aneurysm (AAA) is defined as a localized degeneration of the aortic wall resulting in >50% permanent dilation, hallmarked by progressive inflammatory infiltrate and medial wall degradation.^1, 2^ With over 1 million American adults affected by the disease, the prevalence is ∼2 – 3% in males and 0.5 – 1% of females over the age of 65, resulting in ∼9,928 deaths per year in the US.^3^ Currently, only surgical repair for large (≥55 mm) or rapidly expanding aneurysms are available to prevent aortic rupture, a frequently fatal event.^4^ Understanding the multifactorial complexities of AAA and identifying high risk patients with predictive biomarkers is necessary to improve treatment strategies for aneurysm patients.

A prominent feature of AAA pathogenesis is the formation of a non-occlusive intraluminal thrombus (ILT) driven by increased platelet activity and impaired hemostasis.^5, 6^ While ILT formation may stabilize the aorta during AAA initiation^7^, there is increasing evidence that the ILT contributes to localized inflammation, medial degeneration, and progressive immune infiltrate.^2^ Patients with large and small AAAs have significant increases in circulating hemostatic markers including platelet factor 4 (PF4), thrombin-antithrombin (TAT) complex and D-dimer.^8^ While there are no recognized gold-standard biomarkers for AAA or progression to rupture, the most established is D-dimer, produced as a result of fibrinolysis and indicates active thrombus formation and degradation.^9,10^ A longitudinal study of 299 AAA patients demonstrated D-dimer was significantly associated with AAA growth rate,^11^ while another found D-dimer correlated with sub-aneurysmal dilation.^12^ Conversely, our previous analysis of AAA patients indicated that while D-dimer was associated with AAA incidence, no significant difference was observed between fast (defined as ≥ 2 mm/year) or slow-growing (< 2 mm/year) aneurysms.^8^ While D-dimer could potentially be used as a biomarker for AAA status, its usefulness for predicting AAA progression and rupture remains uncertain.

Platelets are critical to the formation and propagation of the ILT and are largely present at the luminal surface among trapped erythrocytes, neutrophils, and a dense fibrin network.^7, 13^ The platelet-specific receptor glycoprotein VI (GPVI) binds collagen, resulting in platelet activation and adhesion.^14–17^ GPVI interaction with the D-domain of fibrinogen/fibrin augments platelet adhesion and aggregation and facilitates platelet recruitment to an existing clot.^18–21^ Once engaged, GPVI triggers rapid platelet activation via engagement of Immunoreceptor Tyrosine-based Activation Motifs within the noncovalently linked Fc receptor γ-chain dimer, to convey ligand-mediated signaling leading to platelet degranulation.^22, 23^ Activation also triggers autologous proteolysis of GPVI mediated by endogenous ADAM10 in human platelets, liberating a 55-kDa soluble ectodomain fragment known as soluble GPVI (sGPVI).^24, 25^ Constituting the only *bona fide* platelet-specific biomarker of platelet activation,^25^ sGPVI is elevated in prothrombotic conditions including acute ischemic stroke, sepsis, disseminated intravascular coagulation, and deep vein thrombosis.^26^ While traditional antiplatelet therapies may increase bleeding risk in AAA patients,^7, 27–29^ targeting of GPVI in murine models reduces arterial thrombosis without significantly changing bleeding times.^30^ Moreover, patients with a rare homozygous deletion of GPVI present with only mild bleeding diathesis^31, 32^ and genetic deletion of GPVI is not associated with spontaneous bleeding or impaired hemostasis in mice, suggesting GPVI may be a therapeutic target in disorders regulated by activated platelets without affecting hemostasis.^21^ Early phase clinical trials utilizing antibody-mediated inhibition of GPVI have not reported a significant bleeding phenotype.^33–35^ Thus, GPVI is a promising target to reduce platelet-mediated thrombosis whilst avoiding hemorrhagic consequences.

## <u>METHODS</u> (Comprehensive methods found in Supplemental Materials)

### Mouse Study approvals

Mouse studies (protocol #15-01-29-01 or #20-11-05-02) were performed with approval of the University of Cincinnati IACUC.

### Mice

Male low-density lipoprotein receptor deficient (*Ldlr^-/-^*; B6.129S7-*Ldlr^tm1Her^*/J; stock: 2207) and *C57BL/6J* (stock: 000664) mice were obtained from The Jackson Laboratory (Bar Harbor, MA).

### Angiotensin II model of AAA

Male Ldlr-/-(n = 190) mice were infused with sterile saline or AngII (1,000 ng/kg/min) via a subcutaneous osmotic minipump (Alzet Corp) for 28 days, as previously described.^36^

### Topical elastase model of AAA

Male *C57BL/6J* mice (n = 20) underwent laparotomy and a sterile solution of 10mg/mL elastase solution (Millipore Sigma, E7885-5, Lot-SLCL3617) was applied to the infrarenal aorta, as previously described.^37, 38^

### Measurements of abdominal aortic diameters

Abdominal aortas were visualized *in vivo* with ultrasound (Vevo 2100, VisualSonics, Toronto, ON, Canada) as described previously.^39, 40^ Aortas were excised and cleaned before imaging (Nikon SMZ800N) and measuring the maximal outer width (NIS Elements V4.4).^36^

### ELISAs and cytokine array

Human sGPVI was measured using a solid-phase rabbit anti-GPVI polyclonal antibody and fluid-phase murine anti-GPVI monoclonal antibody (1A12) as previously described.^41^

### JAQ1 antibody intervention

At 14 days post-model initiation, mice with established AAAs were blindly randomized and received a one-time 250 µl intraperitoneal injection of JAQ1 (50µg) GPVI neutralizing antibody (Emfret Analytics, Germany) or rat IgG2a (50µg) placebo, as described previously.^30^

### Histological processing of abdominal aortas

Histological processing and staining of aortic sections for CD68 Alexa Fluor® (BioRad, MCA1957-A647) and Picrosirius red (PolyScientific, Inc.) was completed as previously described.^42^

### Human aneurysm samples

Human AAA tissue was obtained from four patients (4 males) aged 65 ± 7.7 years (mean ± SD; range 45 to 71 years) undergoing open aneurysm repair (between 2015 and 2018) at the University of Rochester Medical Center in Rochester, NY under an approved URMC Cardiovascular Tissue Bank IRB (RSRB 00036669). Control tissue was procured from the infrarenal aortas of 5 males aged 58 ± 6.6 years (mean ± SD; range 42 to 65 years) with cause of death deemed non-aortic.

### RNA processing and sequencing

RNA was isolated from human AAA wall, accompanying thrombi, and control aortic wall (described above) with the commercially available NucleoSpin^®^ RNA kit (Macherey-Nagel). RNAseq was performed by the Cincinnati Children’s Gene Expression Core Facility as described previously.^30^

### Human platelet collection

Volunteer control patients (n = 4) with no known medical history, aneurysmal disease, or on antiplatelet agents were recruited in studies approved by the Cleveland Clinic RSRB. Washed AAA patient platelets (n = 5) were collected by venipuncture and isolated from citrate plasma from pre-operative patients under serial imaging surveillance. Platelet collection, immunoblotting, agonist stimulation, and flow cytometry were performed as described previously.^43, 44^

### Research statistics and data representation

Unless indicated otherwise, all bar, scatter, and line graphs were created with Sigma Plot v.14.5 (SPSS, Chicago, IL). Data are represented as individual data points and means ± SEM. Data normality was assessed using a Shapiro Wilk test. Two group comparisons with assumed normality was performed via Student’s t-test. Non-normal data was analyzed with a Mann-Whitney Rank Sum. Multiple group significance was assessed by One-Way ANOVA on Ranks with a Dunn’s post hoc, One Way ANOVA with Holm Sidak Post Hoc, or Two-Way ANOVA with Holm Sidak Post Hoc, when appropriate. Statistical significance among temporal groups was assessed by either a One-Way Repeated Measures (RM) ANOVA (parametric) or RM ANOVA on Ranks (non-parametric), where appropriate. Values of *P* < 0.05 were considered statistically significant.

### European Study Population

We analyzed participants from a population-based case-control study of patients with AAA and healthy controls from the Uppsala region, Sweden, between 2008 and 2016, as described previously and in the online supplemental methods section.^8, 42^

### American Study Population

sGPVI, D-dimer, and AAA were also examined in healthy control or AAA patients from a U.S. case-control study. Details for this cohort have been previously published and are available in the online supplement.^42^

## RESULTS

### Platelet-associated genes are enriched in the ILT of human AAAs

To identify platelet transcripts that may play a role in the pathogenesis of AAA, RNA sequencing was performed on age-matched control aortic tissue (n = 5), AAA aortic tissue (n = 4), and AAA ILT (n = 3). All three groups demonstrate spatial separation via principle component analysis (PCA) (Figure 1A) and heatmap clustering (Figure 1B) with many unique sets of differentially expressed genes in each tissue comparison (Supplemental Figure 1A). Platelet-associated transcripts clustered within the AAA thrombus versus the AAA aortic tissue (Figure 1C). Differential expression analysis of the AAA thrombus versus the AAA wall (Figure 1D) and control aortic wall (Supplemental Figure 1B) demonstrates several platelet transcripts in the upper quartile of significantly enriched genes (noted with red dots) with the volcano plot revealing several significantly upregulated platelet-derived transcripts (Figure 1D – red dots). Platelet transcripts comprise 27% of the top 30 significantly enriched genes in the AAA thrombus vs AAA wall (Figure 1E), including GPVI.

**Figure 1:**
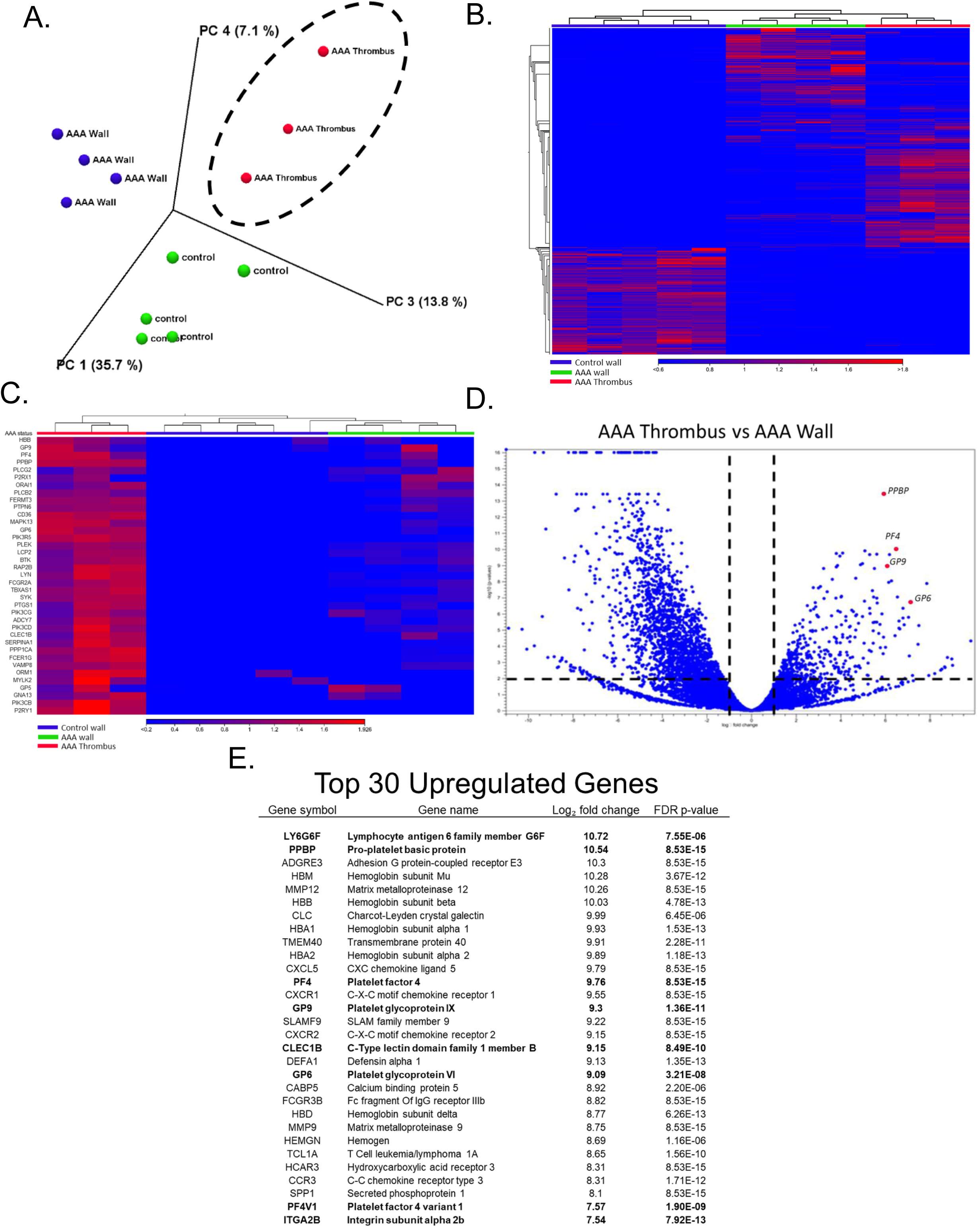
*<u>Platelet-associated gene expression is enriched in human AAA thrombi.</u>* RNA recovered from age-matched control aortic wall (n = 5), AAA aortic wall (n = 4), and AAA thrombi (n = 3) underwent RNA sequencing. (A) Represents the principal component analysis (PCA) of the RNA-seq results from control wall, AAA wall, and AAA thrombus samples. (B) Heat map of differences in gene expression in control wall, AAA wall, and AAA thrombus samples. (C) Heat map of platelet-associated transcripts in AAA thrombus vs. AAA and control aortic tissue. (D) Volcano plot showing total fold change and p-value of significant AAA thrombus genes versus AAA wall, with highlighted platelet-associated genes (red dots). (E) A list of the top 30 significantly enriched genes in AAA thrombus versus AAA wall, platelet-associated genes are bolded.

### The platelet-specific collagen receptor GPVI is upregulated in AAA patients

Analysis of a small cohort of human AAA patients versus age-matched controls demonstrates significantly increased platelet GPVI protein via western blotting and flow cytometry (Figure 2A – 2C). Washed platelets isolated from AAA and control patients were subjected to convulxin, a GPVI agonist. Convulxin-treated platelets from AAA patient had augmented P-selectin expression compared to controls.^45^ This data demonstrates AAA patients have increased GPVI platelet activation.

**Figure 2:**
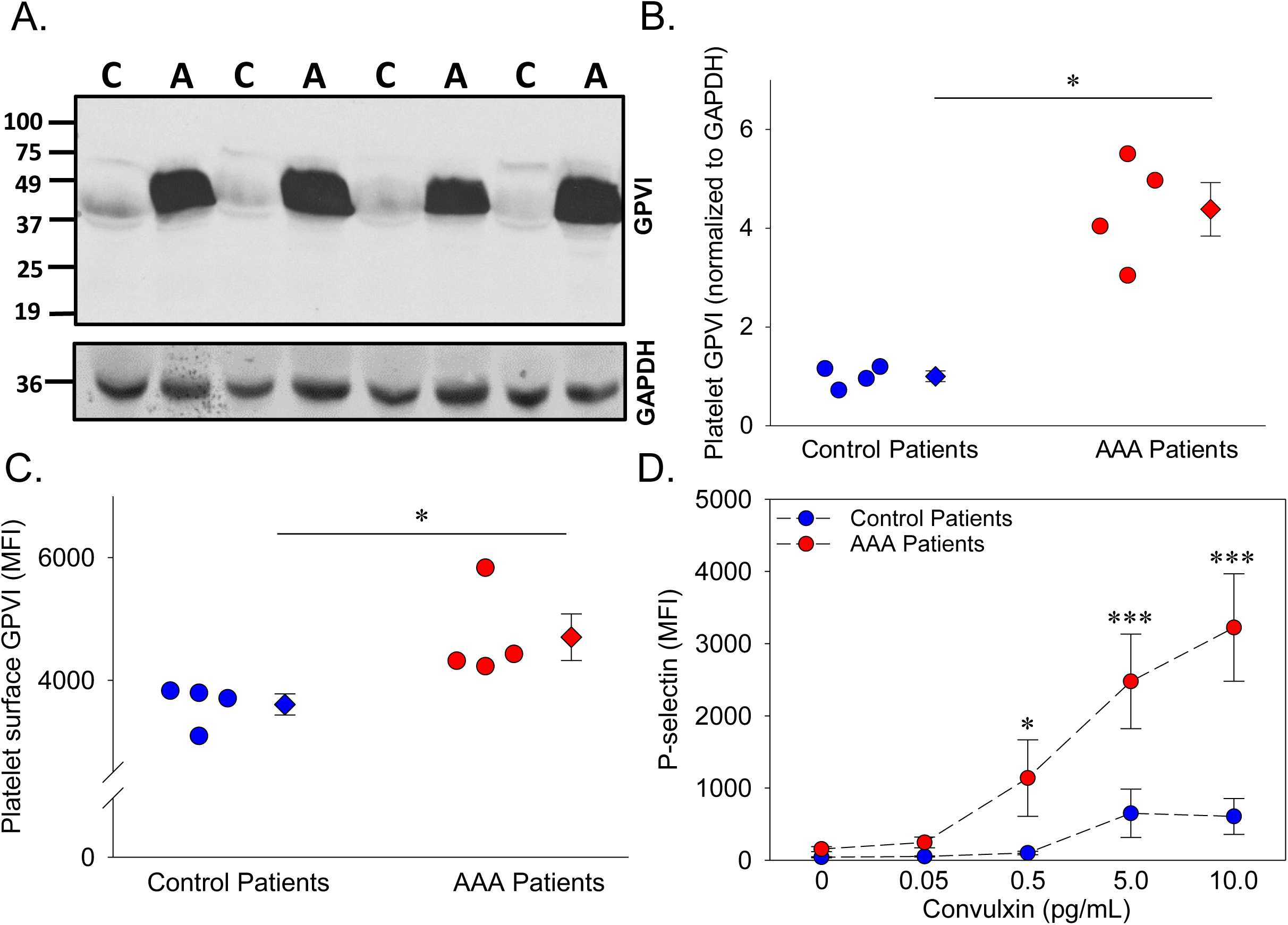
*<u>The thrombotic platelet receptor GPVI is increased in human AAA patients.</u>* Platelets from control (n = 4) and AAA patients (n = 4) were analyzed for the platelet receptor GPVI. Platelet GPVI protein expression is increased in patients with AAA compared to control as measured by Western blot (A) and quantified by densitometry (B) (*, P = 0.007 by Welch’s *t*-test). GPVI on the platelet surface is increased in AAA patients vs. control (C) as demonstrated by platelet surface flow cytometry (*, P = 0.039 by Student’s *t*-test). (D) Platelets isolated from AAA patients demonstrated significantly more surface P-selectin in response to the GPVI specific agonist convulxin compared to control as measured by flow cytometry (*, P = 0.046, ***, P < 0.001 AAA patients vs. control patients by two-way ANOVA). Data represented as individual points or mean ± SEM.

### The platelet marker sGPVI and the fibrinolytic marker D-dimer are elevated in AAA patients and correlate with AAA growth rate

We examined sGPVI concentration in a European cohort of control subjects compared to slow or fast-growing AAAs (Table 1). The predictive value of sGPVI for AAA incidence and growth rate was compared to our published analyses of D-dimer through individual and combined modeling.^8^ Patients with AAA had significantly higher circulating levels of both sGPVI and D-dimer compared to control patients, however, only sGPVI was significantly elevated in fast-growing vs. slow-growing AAA patients (Figure 3A and 3B). Ordinal logistic regression demonstrates the odds of being in a higher outcome group (slow or fast AAA) versus a lower outcome group (control), after adjustment, per doubling of sGPVI was 4.1 (95% CI: 3.0, 5.8). In dichotomous analysis, for participants with sGPVI levels above the cohort median of 38 ng/mL, the odds of being in a higher outcome group was 11.6 (95% CI: 6.2, 21.9) after adjustment (Figure 3C, top). Per doubling of D-dimer, after adjustment the odds of being in the higher outcome group was 2.5 (95% CI: 1.8, 3.6) and 6.1 (95% CI: 3.2, 11.3) in participants with D-dimer levels above the cohort median of 500 ng/mL after adjustment (Figure 3C, bottom).

**Figure 3:**
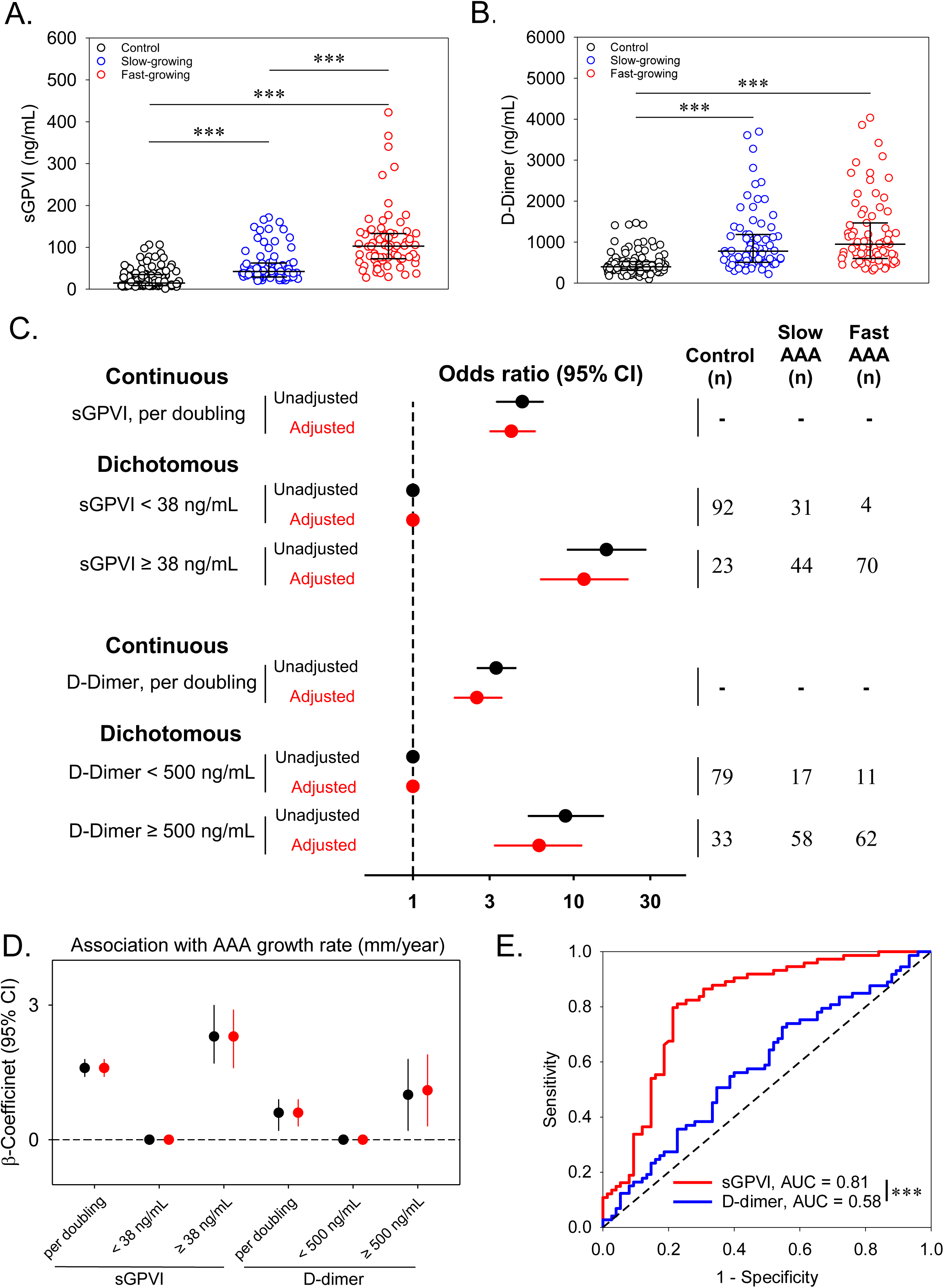
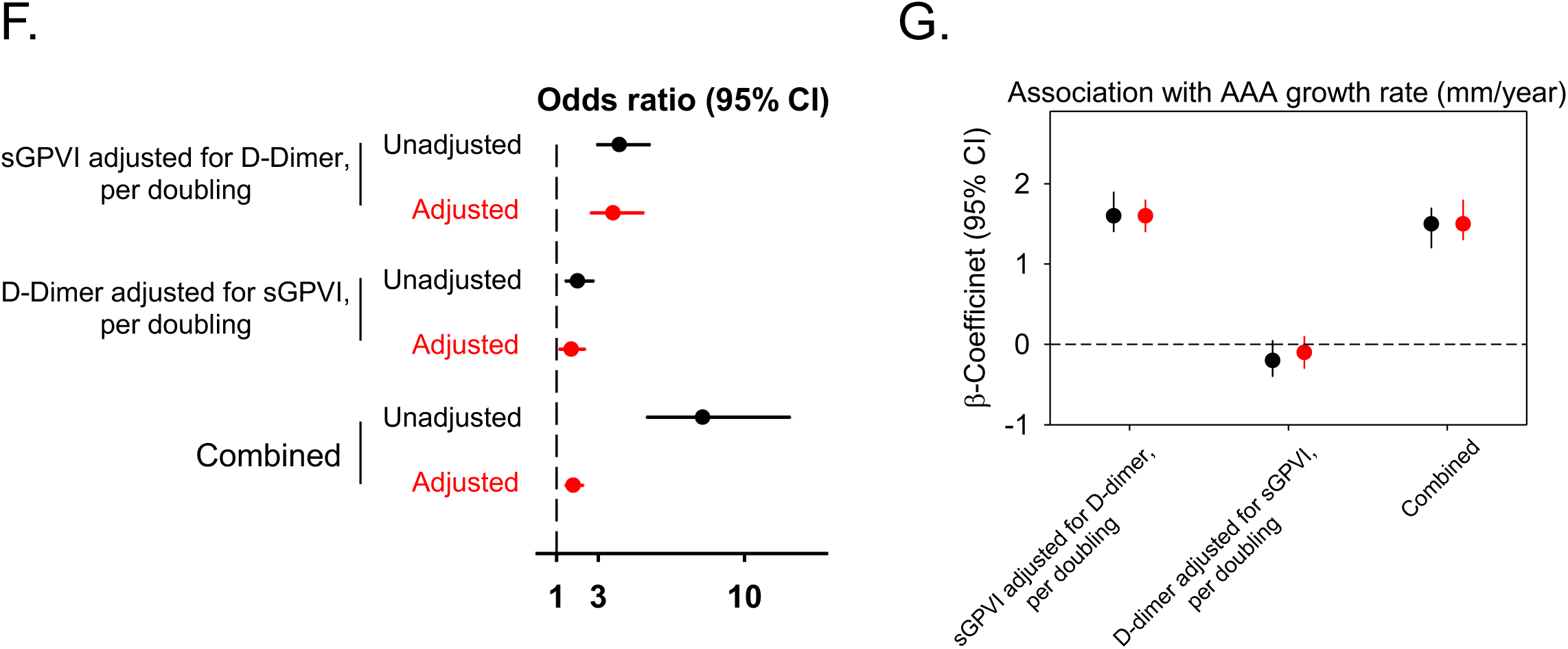
*<u>Circulating sGPVI and D-dimer are increased in AAA patients and are predictive of AAA status and growth rate in the European cohort.</u>* Plasma was assessed for (A) sGPVI and (B) D-dimer from healthy elderly control patients (n = 115), slow-growing AAAs (< 2mm/yr; n = 75), or fast-growing AAAs (> 2mm/yr; n = 74). The median (25^th^, 75^th^ percentile) of sGPVI for controls was 14.9 ng/mL (8.7, 34.9), for slow-growing AAAs was 42.2 ng/mL (29.0, 62.7), and for fast-growing AAAs was 102.9 ng/mL (72.9, 132.9), represented by black bars (***, P < 0.001 by Kruskal-Wallis test). The median (25^th^, 75^th^ percentile) of D-dimer for controls was 397.8 ng/mL (310.0, 525.8), for slow-growing AAAs was 779.5 ng/mL (508.7, 1184.9), and for fast-growing AAAs was 948.1 ng/mL (599.5, 1466.1), represented by black bars (***, P < 0.001 by Kruskal-Wallis test). The OR and 95% CI for the association between (C, top) sGPVI and case status and (C, bottom) D-dimer and case status. Both sGPVI and D-dimer were modeled dichotomously (above/below median) and continuously (log base-2 transformed). (D) Unadjusted (black) and adjusted (red) linear regression for the relationship between sGPVI (n=149), D-dimer (n=148), and the growth rate (mm/year) of AAA. Continuous sGPVI and D-dimer are log transformed, analysis restricted to cases only. (E) Comparative ROC curve analysis of sGPVI and D-dimer to distinguish fast from slow-growing AAA using slow-growing AAAs patients used as reference, (***, P <0.001 as determined by ROC Curve Area Comparison). The effect of (F, top) sGPVI adjusted for D-dimer and (F, middle) D-dimer adjusted for sGPVI on case status was examined using ordinal logistic regression. (F, bottom) The combined effect calculated using linear combination of sGPVI and D-dimer. (G) Unadjusted (black) and adjusted (red) linear regression for the relationship between sGPVI, D-dimer, and the growth rate (mm/year) of AAA. Continuous sGPVI and D-dimer are log transformed, analysis restricted to cases only (n = 148).

**Table 1.** Participant characteristics by case definition (n=264)

| Characteristic | Control<br>(n=115) | Slow-growing<br>(n=75) | Fast-growing<br>(n=74) |
| --- | --- | --- | --- |
| Age (years) | 67.4 (2.6) | 72.7 (6.6) | 72.6 (6.8) |
| Height (cm) | 177.1 (10.3) | 181.8 (6.2) | 180.7 (6.2) |
| Weight (kg) | 84.1 (13.6) | 90.2 (15.7) | 89.3 (16.2) |
| CHD | 10 (8.7) | 27 (36.0) | 25 (33.8) |
| Hypertension | 56 (48.7) | 51 (68.0) | 44 (59.5) |
| CVD | 5 (4.4) | 18 (24.0) | 7 (9.5) |
| Claudication | 1 (0.9) | 8 (10.7) | 1 (1.4) |
| COPD | 5 (4.4) | 12 (16.0) | 11 (14.9) |
| Renal Insufficiency | 2 (1.7) | 4 (5.3) | 4 (5.4) |
| Diabetes | 17 (14.8) | 17 (22.7) | 4 (5.4) |
| Aspirin Use | 17 (14.8) | 37 (49.3) | 33 (44.6) |
| Statin Use | 30 (26.1) | 35 (46.7) | 27 (36.5) |
| Current Smoker | 6 (5.2) | 12 (16.0) | 25 (33.8) |
| Smoking Years | 23.4 (13.7) | 34.2 (13.0) | 39.4 (14.3) |

Note: Values are mean (SD) or N (%); d-dimer is missing in 4 participants
Abbreviations: S.D. standard deviation; sGPVI, soluble glycoprotein VI; cm, centimeters; kg, kilograms; ng, nanograms; mL, milliliters; CHD, coronary heart disease; CVD, cardiovascular disease; COPD, chronic obstructive pulmonary disease,

Linear regression modeling was applied to determine the association between sGPVI, D-dimer, and AAA growth rate. The mean (SD) for AAA growth rate in patients with slow-growing was 0.8 (1.0) mm/year while fast-growing had a growth rate of 3.7 (1.6) mm/year. After adjustment, per doubling of sGPVI, the β-coefficient for growth rate was 1.6 (95% CI: 1.4, 1.8) (Figure 3D). Per doubling of D-dimer, the β-coefficient for growth rate was lower in magnitude at 0.6 (95% CI: 0.3, 0.9) (Figure 3D) after adjustment when compared to sGPVI. Moreover, in a comparative ROC analysis of AAA patients, plasma sGPVI concentration (n = 149) predicts the occurrence of fast-growing AAAs (AUC = 0.81) with an optimum threshold of >64.5 ng/mL with 77% specificity (95% CI: 66.7, 85.3) and 81% sensitivity (95% CI: 70.7, 88.4) significantly better than D-dimer concentrations (n = 148) (AUC = 0.58) with an optimum threshold of >804 ng/mL with 56% specificity (95% CI: 44.8, 66.7) and 58% sensitivity (95% CI: 46.1, 68.2), when slow-growing AAA patients are used as control (Figure 3E).

To further understand the prognostic value of sGPVI and D-dimer, both ordinal logistic and linear regression models were reanalyzed with adjustments for sGPVI and D-dimer. Per doubling of sGPVI, when adjusted for covariates and D-dimer, the odds of being in the higher outcome group was 3.7 (95% CI: 2.6, 5.2) while the OR per doubling of D-dimer, adjusted for covariates and sGPVI, was 1.7 (95% CI: 1.1, 2.4) (Figure 3F). The combined effect of sGPVI and D-dimer on case status was significant after adjustment, with an OR of 1.8 (95% CI 1.4, 2.3) (Figure 3F). After adjusting for comorbidities and D-dimer, per doubling of sGPVI, the β-coefficient for growth rate was significant at 1.6 (95% CI: 1.4, 1.9) while, after adjustment for sGPVI, the association between D-dimer and growth rate was no longer significant (Figure 3G). After adjustment, per doubling of sGPVI and D-dimer combined, the β-coefficient for growth rate was 1.5 (95% CI: 1.3, 1.8) (Figure 3G).

We additionally examined the relationship between sGPVI, D-dimer, and AAA in participants from a separate American case-control study (Table 2). Concordant with results in the European cohort, both plasma sGPVI and D-dimer were significantly increased in AAA patients as compared to control, with only sGPVI levels being significantly increased in fast-growing vs. slow-growing AAA patients (Figure 4A & B). Ordinal logistic regression demonstrates that per doubling of sGPVI, the odds of being in a higher outcome group was 2.4 (95% CI: 1.9, 3.0) and participants with sGPVI levels above the cohort median, 21 ng/mL, had an OR of 5.2 (95% CI: 3.0, 9.2) after adjustment (Figure 4C, top). In contrast to the European cohort, the effect of D-dimer per doubling (OR: 7.0, 95% CI: 4.4, 10.9) was higher in magnitude than the effect of sGPVI. Results were similar in the American cohort for the associations between sGPVI, D-dimer, and growth rate, although in participants with D-dimer levels above 455 ng/mL, the association between D-dimer and growth rate was no longer significant (Figure 4D). Comparative ROC curve analysis using AAA patients (n = 118) demonstrates that sGPVI (AUC = 0.85) with an optimum threshold of >51.8 ng/mL with 83% specificity (95% CI: 74.3, 89.6) and 66% sensitivity (95% CI: 47.4, 80.1) was significantly more predictive of fast-growing AAA compared to D-dimer (AUC = 0.58) with an optimum threshold of >852 ng/mL with 53% specificity (95% CI: 43.1, 63.3) and 79% sensitivity (95% CI: 61.6, 90.2) using slow-growing AAA patients as control (Figure 4E).

**Figure 4:**
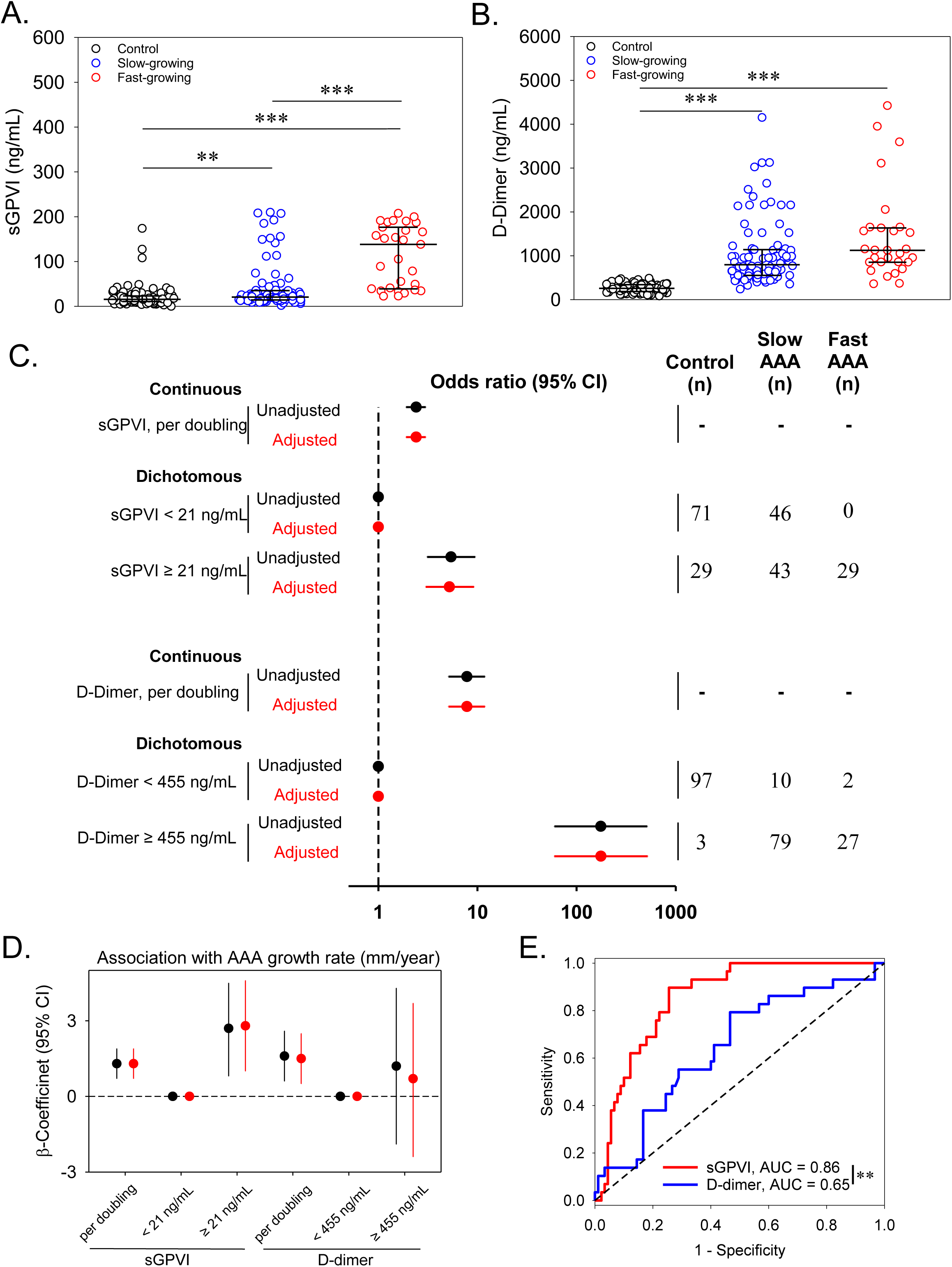
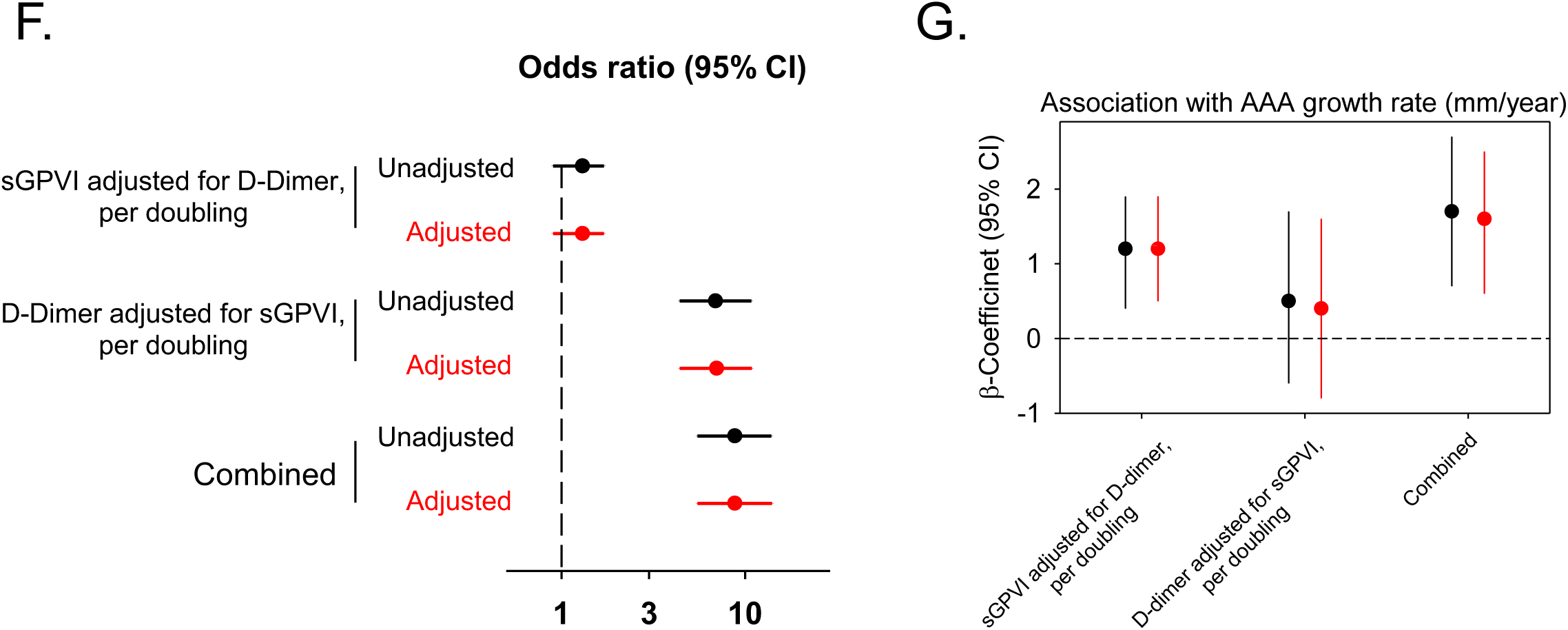
*<u>Circulating sGPVI and D-Dimer are elevated in AAA patients and are predictive for case status and growth rate in the American-case control study.</u>* Plasma was assessed for (A) sGPVI and (B) D-dimer from healthy elderly control patients (n = 100), slow-growing AAAs (< 4mm/yr; n = 89), or fast-growing AAAs (> 4mm/yr; n = 29). The median (25^th^, 75^th^ percentile) of sGPVI for controls was 15.8 ng/mL (10.8, 23.2), for slow-growing AAAs was 20.9 ng/mL (14.3, 35.2), and for fast-growing AAAs was 138.2 ng/mL (39.2, 176.4), represented by black bars (***, P < 0.001 by Kruskal-Wallis test). The median (25^th^, 75^th^ percentile) of D-dimer for controls was 256 ng/mL (184, 340), for slow-growing AAAs was 795 ng/mL (551, 1139), and for fast-growing AAAs was 1123 ng/mL (854, 1633), represented by black bars (**, P = 0.002, ***, P < 0.001 by Kruskal-Wallis test). The OR and 95% CI for the association between (C, top) sGPVI and case status and (C, bottom) D-dimer and case status. Both sGPVI and D-dimer were modeled dichotomously (above/below median) and continuously (log base-2 transformed). (D) Unadjusted (black) and adjusted (red) linear regression for the relationship between sGPVI (n = 118), D-dimer (n = 118), and the growth rate (mm/year) of AAA. Continuous sGPVI and D-dimer are log transformed, analysis restricted to cases only. (E) Comparative ROC analysis using sGPVI and D-dimer to predict the occurrence of fast-growing AAAs using slow-growing AAAs patients used as reference, (***, P = 0.04 as determined by ROC Curve Area Comparison). The effect of (F, top) sGPVI adjusted for D-dimer and (F, middle) D-dimer adjusted for sGPVI on case status was examined using ordinal logistic regression. (F, bottom) The combined effect calculated using linear combination of sGPVI and D-dimer. (G) Unadjusted (black) and adjusted (red) linear regression for the relationship between sGPVI, D-dimer, and the growth rate (mm/year) of AAA. Continuous sGPVI and D-dimer are log transformed, analysis restricted to cases only (n = 118).

**Table 2.**
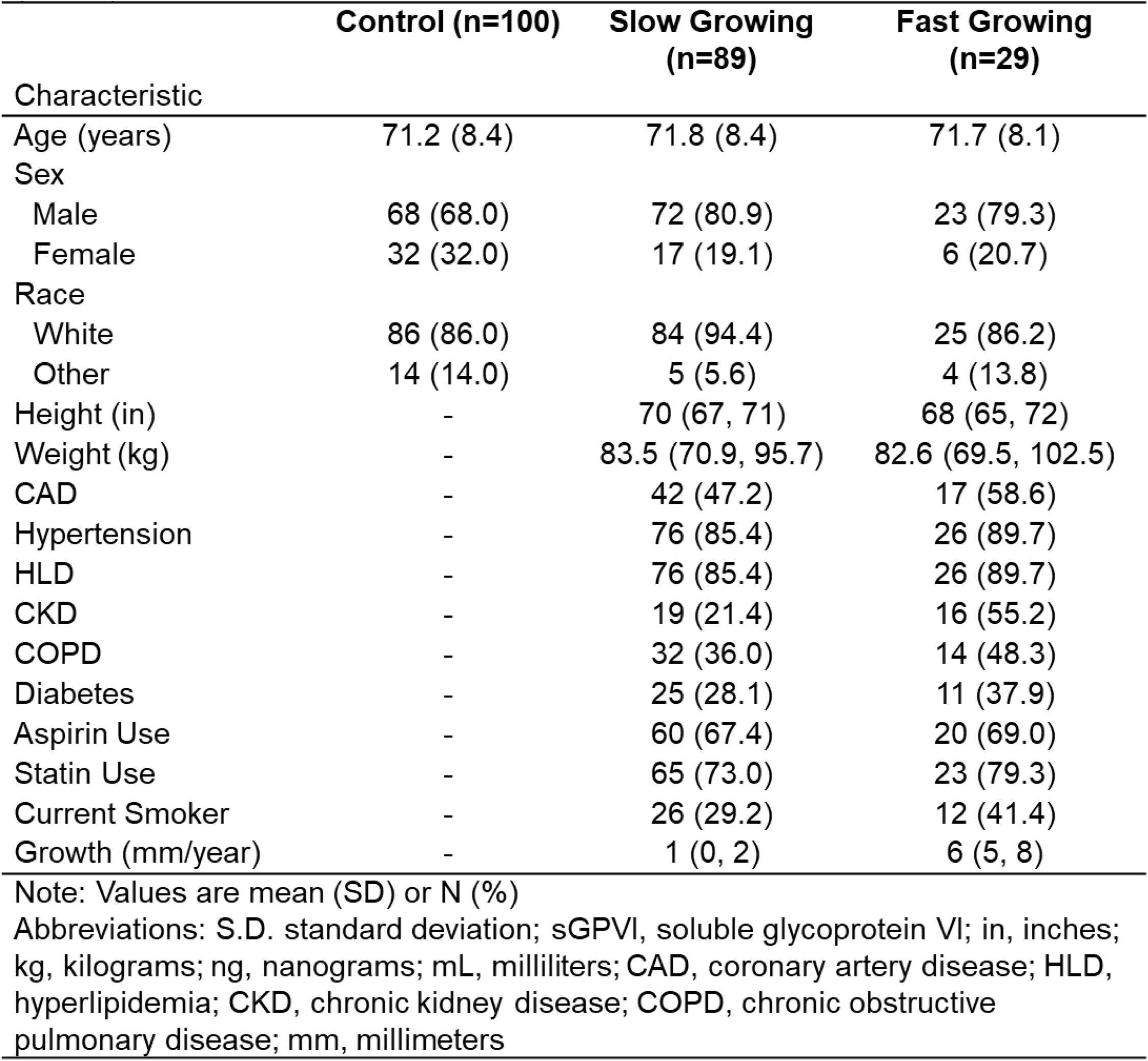
Participant characteristics of the validation study by case definition (n=218)

The combined effects of sGPVI and D-dimer on case status and growth rate for the American case control study are presented in Figure 4F. In contrast with results in the European cohort, the effect of D-dimer (OR: 7.0, 95% CI: 4.4, 10.9) was higher in magnitude than the effect of sGPVI (OR: 1.3, 95% CI: 0.9, 1.7) on case status in the American cohort. Interestingly, after adjustment for sGPVI, D-dimer was not associated with growth rate (β-coefficient: 0.4, 95% CI: -0.8, 1.6) while sGPVI, after adjustment D-dimer, had a significant association with growth rate (β-coefficient: 1.2, 95% CI: 0.5, 1.9) (Figure 4G). Taken together, these results suggest that both sGPVI and D-dimer can assist in AAA diagnosis, while sGPVI is more informative with respect to predicting an accelerated rate of AAA growth.

### Platelet targeting via GPVI Inhibition via JAQ1 antibody intervention blunts progression of established AAAs in murine models

In mice, JAQ1 antibody injection inhibits GPVI-mediated platelet activation with minimal effect on bleeding times.^30^ To test the efficacy of targeting GPVI to reduce AAA burden, male *Ldlr^−/−^* underwent AngII-infusion for 28 days. At day 14, mice were blindly randomized into two groups normalized for aortic diameter, receiving either a one-time IP injection of JAQ1 antibody or rat IgG2a control (50μg; Figure 5A). JAQ1 antibody injection did not affect body weight (Supplemental Figure 2A). Mice that received JAQ1 antibody demonstrated significantly less aortic growth post intervention compared to control treated mice (Figure 5B). Moreover, JAQ1 treated mice experienced zero rupture-induced deaths while 3/9 mice in the control group died from aortic rupture (Supplemental Figure 2B). Final aortic diameter was assessed *ex vivo* with JAQ1 treated mice having significantly smaller aortic diameters compared control mice (Figure 5C & D). Additionally, J-treated mice had significantly more Type I collagen, as measured by picrosirius red (Supplemental Figure 2C & D), and less CD68 infiltrating macrophages (Supplemental Figure 2C & E) compared to control treated mice.

**Figure 5:**
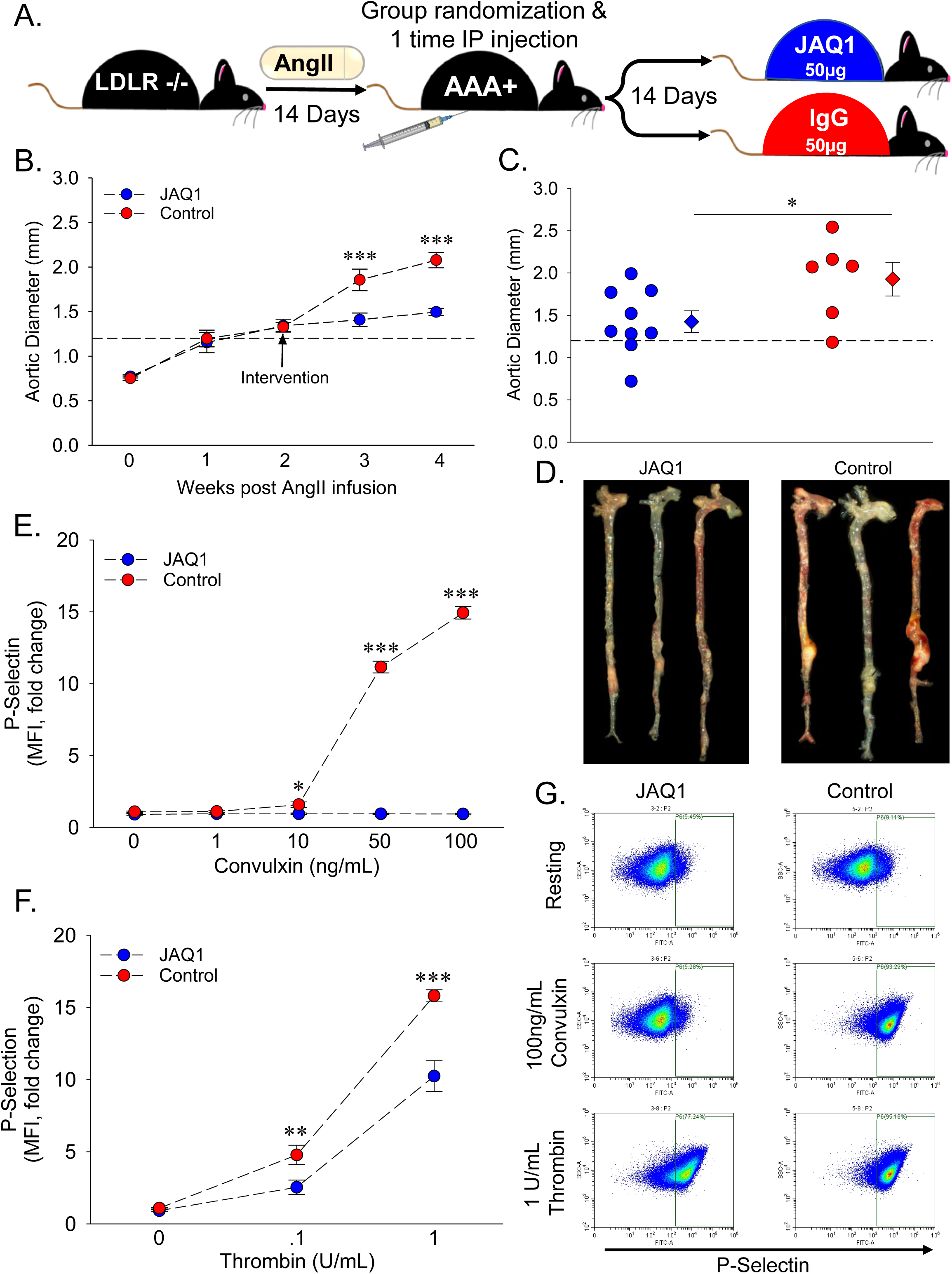
*<u>GPVI blockade via intervention with JAQ1 antibody blunts AngII-induced AAA expansion and rupture-induced death.</u>* (A) Study schematic. Male *Ldlr^-/-^* mice (8 – 10 weeks old) were infused with angiotensin II (AngII; 1,000 ng/kg/day) via osmotic pump implantation. After 14 days of AngII-infusion, mice were blindly randomized into two groups of controlled for aortic diameter and given either a one-time IP injection of 50µg JAQ1 antibody (n=9) or 50µg IgG control (n=9). Following another 14 days, mice were euthanized, final aortic diameter was measured *ex vivo*, and tissues were harvested for further analysis. (B) Intraluminal aortic diameters measured by weekly ultrasound (***, P < 0.001 JAQ1 vs. Control by two-way repeated measures ANOVA). (C) Aortic diameters in JAQ1 treated vs. control treated mice measured *ex vivo* 28 days after AngII-infusion (*, P = 0.045 by Student’s t-test). (D) Representative images of aortas *ex vivo* from the indicated groups. Platelet activation measured 14 days post JAQ1 or IgG antibody intervention indicated by surface P-selectin (FITC) levels in response to varying doses of the GPVI specific agonist (E) convulxin or (F) thrombin in JAQ1 treated vs, control mice measured flow cytometry (n = 6/group. *, P = 0.032, **, P = 0.01, ***, P < 0.001 JAQ1 vs. Control by two-way ANOVA). (G) Representative flow cytometry scatter plots from the indicated conditions. Data represented as individual data points or mean ± SEM.

To confirm that JAQ1 treatment resulted in GPVI blockade, washed platelets were prepared from JAQ1 (n = 6) and control (n=6) treated mice 28 days post AngII-infusion. Control platelets activated in a dose dependent manner to both convulxin (GPVI agonist) or thrombin (murine platelet PAR3 and PAR4 agonist) ^46^ stimulation (Figure 5E & F). While platelets from J-treated mice partially responded to thrombin, convulxin treatment elicited no activation, indicating successful and highly specific blockade of the platelet GPVI receptor (Figure 5E & F). Interestingly, JAQ1 treated mice had significantly lower activation in response to thrombin compared to control (Figure 5F). There was no difference in platelet count between groups (Supplemental Figure 2F).

As a second model of AAA, male *C57BL/6J* mice were subjected to open laparotomy and topical elastase application (10mg/mL for 5 minutes) to the infrarenal aorta. Blinded randomization into JAQ1 (n = 10) and control (n = 9) intervention groups with equal aortic diameter was performed on day 14 (Figure 6A). Similar to the AngII model, mice treated with JAQ1 had significantly lower aortic diameter after intervention compared to control (Figure 6B, C, & D). JAQ1 treated mice had significantly more type I collagen (Supplemental Figure 3C left & D), however no difference in macrophage accumulation was detected (Supplemental Figure 3C right & E). Moreover, treatment of washed platelets with convulxin and thrombin demonstrates that JAQ1 intervention specifically inhibited GPVI-mediated platelet activation (Figure 6E, F, & G). The congruent results obtained in two murine AAA models suggest platelet inhibition by selective GPVI targeting is a feasible therapeutic option to limit AAA progression.

**Figure 6:**
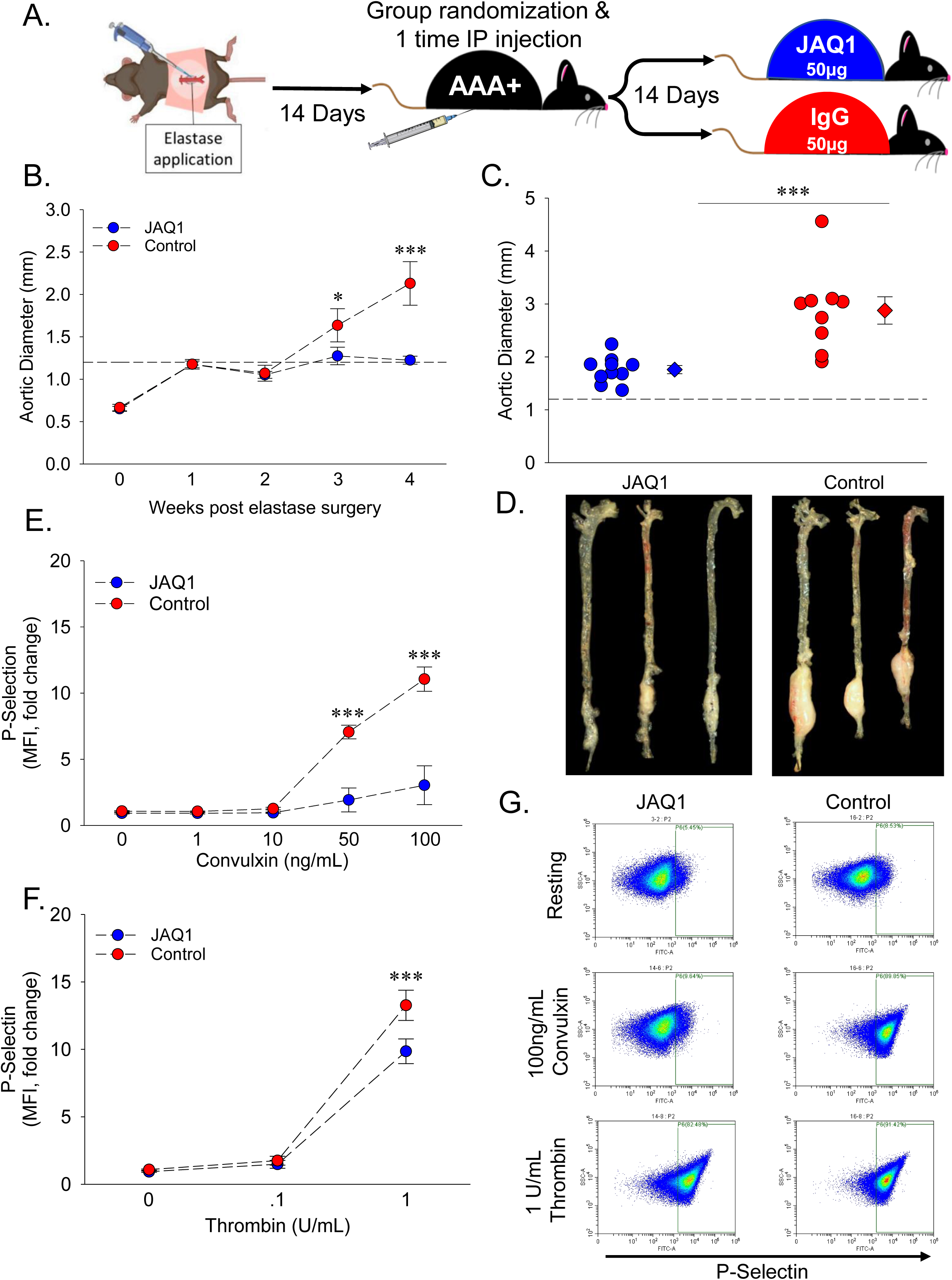
*<u>GPVI blockade via intervention with JAQ1 antibody mitigates Elastase AAA expansion.</u>* (A) Study schematic. Male *Ldlr^-/-^* mice (8 – 10 weeks old) underwent laparotomy and elastase (5μl of 10 mg/mL porcine pancreatic elastase) was applied to the adventitia of infrarenal aorta for 5 minutes to initiate AAA. At 14 days post elastase surgery, mice were blindly randomized into two groups controlled for aortic diameter and given either a one-time IP injection of 50µg JAQ1 antibody (n = 10) or 50µg IgG control (n = 9). Following another 14 days, mice were euthanized, final aortic diameter was measured *ex vivo*, and tissues were harvested for further analysis. (B) Intraluminal aortic diameters measured by weekly ultrasound (*, P = 0.026, ***, P < 0.001 JAQ1 vs. Control by two-way repeated measures ANOVA). (C) Aortic diameters in JAQ1 treated vs. control treated mice measured *ex vivo* 28 days after elastase surgery (***, P < 0.001 by Rank Sum Test). (D) Representative images of aortas *ex vivo* from the indicated groups. (D) Representative images of aortas *ex vivo* from the indicated groups. Platelet activation measured 14 days post JAQ1 or IgG antibody intervention indicated by surface P-selectin (FITC) levels in response to varying doses of the GPVI specific agonist (E) convulxin or (F) thrombin in JAQ1 treated vs, control mice measured flow cytometry (n = 6/group. ***, P < 0.001 JAQ1 vs. Control by two-way ANOVA). (G) Representative flow cytometry scatter plots from the indicated conditions. Data represented as individual data points or mean ± SEM.

## DISCUSSION

This study, in both humans with AAA and in two independent murine models of AAA, revealed GPVI as the first potential target offering *both* diagnostic and therapeutic potential in patients with infrarenal aortic aneurysmal disease. The latest guidelines issued jointly by American Heart Association and the American College of Cardiology give the platelet inhibitor aspirin a Class IIb recommendation in managing patients with AAA with ILT or penetrating aortic ulcer, but also with the presumption of concomitant arterial atherosclerosis.^47^ However, this recommendation was made by expert consensus rather than the availability of mechanistic data.^47^ Our data demonstrates that GPVI is enriched in AAA-associated ILTs, functionally increased in AAA patients, and that sGPVI is positively associated with an aneurysmal infrarenal aorta as well as aortic aneurysm growth rate. This observation was fully reproducible in two independent human cohorts evaluated in two different countries. Additionally, comparing sGPVI with D-dimer (the strongest biomarker to date for predicting adverse AAA outcomes), sGPVI was demonstrated to better predict AAA progression in patients with established infrarenal aortic aneurysmal disease. Taken together, our data indicates that while D-dimer and sGPVI similarly predict a diagnosis of AAA, blood sGPVI concentration more easily identifies patients with fast AAA growth and reveals itself to be a very useful biomarker for clinical decision-making.

The size and growth rate of an ILT directly correlates with risk of AAA rupture and exists in a balance of active thrombus generation and resolution, although total resolution is rarely achieved.^13, 48^ As the most robust blood biomarker of platelet activation, elevated sGPVI suggests platelet adhesion and deposition at the luminal surface of the ILT, which is a dynamic and progressive process that may promote AAA expansion. Conversely, D-dimer is a product of clot resolution and while levels in circulation are highly sensitive for detecting intravascular coagulation and clot resolution, it is not a direct measure of thrombus formation.^49^ While D-dimer undoubtedly indicates the presence of ILT associated with AAA, our data indicates that sGPVI is a more sensitive marker of ongoing ILT formation and active AAA growth with potential as a biomarker for the determination of surgical intervention.

A strength of the present study is the use of two independent patient cohorts from distinct geographic regions which favors generalizability of the scientific observations and allows us to compare separate populations with different modifiable risk factors. While hypertension and smoker status were prevalent in the European cohort, the American cohort had more comorbidities including COPD and diabetes. Additionally, the American cohort included women (Table 2), a population with a lower incidence of AAA, but an increased preponderance for unpredictable ruptures occurring at smaller abdominal diameters.^4, 50, 51^ Control patient covariates including height, weight, smoking, comorbidities, medication use, and aorta diameter were not available for the American cohort which is a limitation. Female subjects for the European cohort were excluded as no appropriate controls were available. Another strength is the use of a population-based case-control study design for both cohorts and the relatively large sample size of slow-and fast-growing AAA in the European cohort. Using both cohorts, we were able to reanalyze and directly compare our previously published results on the association of D-dimer,^8^ the most established biomarker for AAA, to our novel analysis of sGPVI. While aortic diameter was assessed longitudinally in both cohorts, no further follow up data was available regarding rupture or surgical intervention as these patients were excluded. Subsequently, further prospective trials including longitudinal plasma sampling are needed to fully define parameters for D-dimer and sGPVI as biomarkers of AAA status and growth rate.

We have identified a role for platelet activation in the progression of vascular degeneration and rupture in a mouse model of AAA.^2, 52^ Recent work from our collective group demonstrates a mechanistic link between increased platelet reactivity and AAA progression with concomitant increases in platelet receptor expression.^53^ This suggests that the circulating platelet in AAA has a divergent phenotype from healthy conditions and may offer diagnostic as well as therapeutic potential. Studies targeting platelets with GPIIb/IIIa or P2Y_12_ inhibitors successfully decreased AAA burden in rodent models^2, 54, 55^, however, results from clinical data with respect to antiplatelet drugs’ effects on AAA outcome in humans is mixed.^7, 29^ Current platelet inhibitors are associated with bleeding, limiting their utility in AAA patients, creating a need for the development of alternative anti-platelet targets. GPVI antagonism provides a favorable approach to limit pathologic platelet aggregation without bleeding.^21, 30, 56–59^ The minimal bleeding phenotype associated with loss of GPVI is attributed to the existence of several redundant collagen-responsive platelet activation pathways including integrin α2β1 and GPV receptors, as well as GPIbα and αIIβ3 interaction with collagen-bound von Willebrand factor.^21, 60–62^ While this suggests that platelet-mediated primary hemostasis is not dependent on GPVI activity, GPVI is critical to platelet adhesion, propagation, and stability of an existing thrombus via efficient phosphorylation of secondary messengers, robust release of Ca^2+^ and ADP, and increased activity of integrins in response to collagen binding.^18–21, 63, 64^ Moreover, GPVI binding of fibrin facilitates platelet activation, stable platelet adhesion, and platelet spreading with concomitant augmentation of thrombin generation via increased phosphatidylserine exposure.^18, 21, 65^ The accumulation of activated platelets at the luminal surface of an aneurysm contributes to deposition of new ILT layers and overall AAA growth rate, which has been linked to fatal aortic rupture.^5, 13^ Therefore, we postulate that blockade of GPVI effectively reduces platelet aggregation on the exposed collagen and fibrin matrix present at the site of ILT and in turn blunts AAA progression.

Due to the appeal of targeting platelets to treat cardiovascular disease without affecting primary hemostasis, antagonists of GPVI have been developed and brought into clinical trials. Revacept is a recombinant competitive inhibitory protein ligand for collagen binding to GPVI, thus reducing platelet activation, as seen in experimental models of thrombosis.^66, 67^ Phase II clinical trials show Revacept reduces ischemic lesions in symptomatic internal carotid artery stenosis patients,^68^ however, myocardial injury was unchanged between Revacept and placebo in stable ischemic heart disease (IHD) patients undergoing percutaneous intervention.^69^ While exposed collagen is abundant at the luminal site of AAA, it is structurally abnormal which may limit the effectiveness of Revacept in patients with AAA.^70^ A second clinically-relevant compound is Glenzocimab, which directly binds to GPVI, blocking the interaction with collagen and fibrinogen/fibrin and decreasing platelet adhesion and thrombosis. Glenzocimab combined with r-tPA attenuated symptomatic intracerebral hemorrhage, a condition associated with altered GPVI activity and expression, and improves the survival of patients.^71^ Additionally, Glenzocimab treatment is associated with thrombus disintegration, which could aid in the regression of AAA-associated ILTs and improve patient outcomes. Conversely, destabilization of a large ILT could cause thromboembolism and increase the risk of MI and stroke, as noted in EVAR patients.^72^ Importantly, increased bleeding times have not been observed with Revacept or Glenzocimab and both are well-tolerated by patients.^35, 65^ Yet, these compounds have not been tested in AAA patients. Given that the risk of adverse bleeding events and AAA incidence increases with age, it is vital to develop therapeutics that reduce AAA burden without bleeding side effects. Targeting GPVI with antagonists in clinical development presents an exciting opportunity as the first viable therapeutic intervention for AAA patients, but more work is needed to fully understand the implications of such a strategy in aneurysmal pathogenesis.

## Acknowledgements

This work was supported by National Institutes of Health grants R01-HL147171 (APOIII); R01-HL158801; T32 HL125204 (TWB); Swedish Research Council grant K2013-64X-20406-07-3 and grant K2013-99X-22275-01-3 (AW and MB), the Swedish Heart-Lung Foundation grants 2012-0353 and 2015-0596 (AW and MB), and the Konung Gustaf V:s och Drottning Victorias Frimurarestiftelse (AW); Clinical Translational Science Award TL1TR002244 from the National Center for Advancing Translational Sciences (MMP). Aspects of the project were supported by the National Center for Advancing Translational Sciences of the National Institutes of Health, under Award Number 2UL1TR001425.

## Authorship Contributions

T.W.B., A.S., H.M.R., S.J.C, N.M., and A.P.O. conceived and designed the study. T.W.B., A.S., H.M.R., T.M.C, S.I.B., and C.W-L. were responsible for animal care. S.J.C., R.B., S.S., D.M., S.L., A.A., M.P. were responsible for phlebotomy, clinical data management, and regulation. M.M.P. performed statistical analysis of human cohort data. T.W.B., S.J.C., A.A., R.B., A.S., H.M.R., T.M.C, S.I.B., C.W-L., and A.P.O. conducted laboratory testing. T.W.B., A.S., H.M.R., M.M.P., S.S., M.P. and A.P.O. contributed to data processing. T.W.B., A.S., and A.P.O. supervised all aspects of the study. T.W.B., A.S., M.M.P., and A.P.O., contributed to initial data interpretation and wrote the manuscript. All authors contributed to final data interpretation and critical revision of the manuscript and approved the final version of the manuscript.

## REFERENCES

1. Daugherty A, Cassis LA. Mouse models of abdominal aortic aneurysms. Arterioscler Thromb Vasc Biol. 2004;24:429–434. doi: 10.1161/01.ATV.0000118013.72016.ea

2. Owens AP, 3rd, Edwards TL, Antoniak S, Geddings JE, Jahangir E, Wei WQ, Denny JC, Boulaftali Y, Bergmeier W, Daugherty A, et al. Platelet Inhibitors Reduce Rupture in a Mouse Model of Established Abdominal Aortic Aneurysm. Arterioscler Thromb Vasc Biol. 2015;35:2032–2041. doi: 10.1161/atvbaha.115.305537

3. Summers KL, Kerut EK, Sheahan CM, Sheahan MG, 3rd. Evaluating the prevalence of abdominal aortic aneurysms in the United States through a national screening database. J Vasc Surg. 2021;73:61–68. doi: 10.1016/j.jvs.2020.03.046

4. Chaikof EL, Dalman RL, Eskandari MK, Jackson BM, Lee WA, Mansour MA, Mastracci TM, Mell M, Murad MH, Nguyen LL, et al. The Society for Vascular Surgery practice guidelines on the care of patients with an abdominal aortic aneurysm. J Vasc Surg. 2018;67:2–77.e72. doi: 10.1016/j.jvs.2017.10.044

5. Stenbaek J, Kalin B, Swedenborg J. Growth of thrombus may be a better predictor of rupture than diameter in patients with abdominal aortic aneurysms. Eur J Vasc Endovasc Surg. 2000;20:466–469. doi: 10.1053/ejvs.2000.1217

6. Scott DJ, Prasad P, Philippou H, Rashid ST, Sohrabi S, Whalley D, Kordowicz A, Tang Q, West RM, Johnson A, et al. Clot architecture is altered in abdominal aortic aneurysms and correlates with aneurysm size. Arterioscler Thromb Vasc Biol. 2011;31:3004–3010. doi: 10.1161/atvbaha.111.236786

7. Cameron SJ, Russell HM, Owens AP, 3rd. Antithrombotic therapy in abdominal aortic aneurysm: beneficial or detrimental? Blood. 2018;132:2619–2628. doi: 10.1182/blood-2017-08-743237

8. Sundermann AC, Saum K, Conrad KA, Russell HM, Edwards TL, Mani K, Bjorck M, Wanhainen A, Owens AP, 3rd. Prognostic value of D-dimer and markers of coagulation for stratification of abdominal aortic aneurysm growth. Blood Adv. 2018;2:3088–3096. doi: 10.1182/bloodadvances.2017013359

9. Weitz JI, Fredenburgh JC, Eikelboom JW. A Test in Context: D-Dimer. J Am Coll Cardiol. 2017;70:2411–2420. doi: 10.1016/j.jacc.2017.09.024

10. Davies RS, Abdelhamid M, Wall ML, Vohra RK, Bradbury AW, Adam DJ. Coagulation, fibrinolysis, and platelet activation in patients undergoing open and endovascular repair of abdominal aortic aneurysm. J Vasc Surg. 2011;54:865–878. doi: 10.1016/j.jvs.2011.04.010

11. Golledge J, Muller R, Clancy P, McCann M, Norman PE. Evaluation of the diagnostic and prognostic value of plasma D-dimer for abdominal aortic aneurysm. Eur Heart J. 2011;32:354–364. doi: 10.1093/eurheartj/ehq171

12. Vele E, Kurtcehajic A, Zerem E, Maskovic J, Alibegovic E, Hujdurovic A. Plasma D-dimer as a predictor of the progression of abdominal aortic aneurysm. J Thromb Haemost. 2016;14:2298–2303. doi: 10.1111/jth.13487

13. Wilson JS, Virag L, Di Achille P, Karsaj I, Humphrey JD. Biochemomechanics of intraluminal thrombus in abdominal aortic aneurysms. J Biomech Eng. 2013;135:021011. doi: 10.1115/1.4023437

14. Nieswandt B, Watson SP. Platelet-collagen interaction: is GPVI the central receptor? Blood. 2003;102:449–461. doi: 10.1182/blood-2002-12-3882

15. Farndale RW, Sixma JJ, Barnes MJ, de Groot PG. The role of collagen in thrombosis and hemostasis. J Thromb Haemost. 2004;2:561–573. doi: 10.1111/j.1538-7836.2004.00665.x

16. Jandrot-Perrus M, Busfield S, Lagrue AH, Xiong X, Debili N, Chickering T, Le Couedic JP, Goodearl A, Dussault B, Fraser C, et al. Cloning, characterization, and functional studies of human and mouse glycoprotein VI: a platelet-specific collagen receptor from the immunoglobulin superfamily. Blood. 2000;96:1798–1807.

17. Diaz-Ricart M, Tandon NN, Carretero M, Ordinas A, Bastida E, Jamieson GA. Platelets lacking functional CD36 (glycoprotein IV) show reduced adhesion to collagen in flowing whole blood. Blood. 1993;82:491–496.

18. Mammadova-Bach E, Ollivier V, Loyau S, Schaff M, Dumont B, Favier R, Freyburger G, Latger-Cannard V, Nieswandt B, Gachet C, et al. Platelet glycoprotein VI binds to polymerized fibrin and promotes thrombin generation. Blood. 2015;126:683–691. doi: 10.1182/blood-2015-02-629717

19. Mangin PH, Onselaer MB, Receveur N, Le Lay N, Hardy AT, Wilson C, Sanchez X, Loyau S, Dupuis A, Babar AK, et al. Immobilized fibrinogen activates human platelets through glycoprotein VI. Haematologica. 2018;103:898–907. doi: 10.3324/haematol.2017.182972

20. Induruwa I, Moroi M, Bonna A, Malcor JD, Howes JM, Warburton EA, Farndale RW, Jung SM. Platelet collagen receptor Glycoprotein VI-dimer recognizes fibrinogen and fibrin through their D-domains, contributing to platelet adhesion and activation during thrombus formation. J Thromb Haemost. 2018;16:389–404. doi: 10.1111/jth.13919

21. Mangin PH, Gardiner EE, Ariëns RAS, Jandrot-Perrus M. GPVI interplay with fibrin(ogen) in thrombosis. J Thromb Haemost. 2023. doi: 10.1016/j.jtha.2023.03.022

22. Gardiner EE, Arthur JF, Kahn ML, Berndt MC, Andrews RK. Regulation of platelet membrane levels of glycoprotein VI by a platelet-derived metalloproteinase. Blood. 2004;104:3611–3617. doi: 10.1182/blood-2004-04-1549

23. Gardiner EE, Karunakaran D, Arthur JF, Mu FT, Powell MS, Baker RI, Hogarth PM, Kahn ML, Andrews RK, Berndt MC. Dual ITAM-mediated proteolytic pathways for irreversible inactivation of platelet receptors: de-ITAM-izing FcgammaRIIa. Blood. 2008;111:165–174. doi: 10.1182/blood-2007-04-086983

24. Gardiner EE. Proteolytic processing of platelet receptors. Res Pract Thromb Haemost. 2018;2:240–250. doi: 10.1002/rth2.12096

25. Montague SJ, Andrews RK, Gardiner EE. Mechanisms of receptor shedding in platelets. Blood. 2018;132:2535–2545. doi: 10.1182/blood-2018-03-742668

26. Perrella G, Nagy M, Watson SP, Heemskerk JWM. Platelet GPVI (Glycoprotein VI) and Thrombotic Complications in the Venous System. Arterioscler Thromb Vasc Biol. 2021;41:2681–2692. doi: 10.1161/atvbaha.121.316108

27. De Rango P, Verzini F, Parlani G, Cieri E, Simonte G, Farchioni L, Isernia G, Cao P. Safety of chronic anticoagulation therapy after endovascular abdominal aneurysm repair (EVAR). Eur J Vasc Endovasc Surg. 2014;47:296–303. doi: 10.1016/j.ejvs.2013.12.009

28. Rose J, Evans C, Barleben A, Bandyk D, Wilson SE, Chang DC, Lane J. Comparative safety of endovascular aortic aneurysm repair over open repair using patient safety indicators during adoption. JAMA Surg. 2014;149:926–932. doi: 10.1001/jamasurg.2014.1018

29. Wanhainen A, Mani K, Kullberg J, Svensjö S, Bersztel A, Karlsson L, Holst J, Gottsäter A, Linné A, Gillgren P, et al. The effect of ticagrelor on growth of small abdominal aortic aneurysms-a randomized controlled trial. Cardiovasc Res. 2020;116:450–456. doi: 10.1093/cvr/cvz133

30. Nieswandt B, Schulte V, Bergmeier W, Mokhtari-Nejad R, Rackebrandt K, Cazenave JP, Ohlmann P, Gachet C, Zirngibl H. Long-term antithrombotic protection by in vivo depletion of platelet glycoprotein VI in mice. J Exp Med. 2001;193:459–469. doi: 10.1084/jem.193.4.459

31. Nagy M, Perrella G, Dalby A, Becerra MF, Garcia Quintanilla L, Pike JA, Morgan NV, Gardiner EE, Heemskerk JWM, Azócar L, et al. Flow studies on human GPVI-deficient blood under coagulating and noncoagulating conditions. Blood Adv. 2020;4:2953–2961. doi: 10.1182/bloodadvances.2020001761

32. Matus V, Valenzuela G, Sáez CG, Hidalgo P, Lagos M, Aranda E, Panes O, Pereira J, Pillois X, Nurden AT, et al. An adenine insertion in exon 6 of human GP6 generates a truncated protein associated with a bleeding disorder in four Chilean families. J Thromb Haemost. 2013;11:1751–1759. doi: 10.1111/jth.12334

33. Renaud L, Lebozec K, Voors-Pette C, Dogterom P, Billiald P, Jandrot Perrus M, Pletan Y, Machacek M. Population Pharmacokinetic/Pharmacodynamic Modeling of Glenzocimab (ACT017) a Glycoprotein VI Inhibitor of Collagen-Induced Platelet Aggregation. J Clin Pharmacol. 2020;60:1198–1208. doi: 10.1002/jcph.1616

34. Billiald P, Slater AS, Welin M, Clark JC, Loyau S, Pugnière M, Jiacomini IG, Rose N, Lebozec K, Toledano E, et al. Targeting platelet GPVI with glenzocimab: a novel mechanism for inhibition. Blood Adv. 2022. doi: 10.1182/bloodadvances.2022007863

35. Voors-Pette C, Lebozec K, Dogterom P, Jullien L, Billiald P, Ferlan P, Renaud L, Favre-Bulle O, Avenard G, Machacek M, et al. Safety and Tolerability, Pharmacokinetics, and Pharmacodynamics of ACT017, an Antiplatelet GPVI (Glycoprotein VI) Fab. Arterioscler Thromb Vasc Biol. 2019;39:956–964. doi: 10.1161/atvbaha.118.312314

36. Daugherty A, Manning MW, Cassis LA. Angiotensin II promotes atherosclerotic lesions and aneurysms in apolipoprotein E-deficient mice. J Clin Invest. 2000;105:1605–1612. doi: 10.1172/JCI7818

37. Lu G, Su G, Davis JP, Schaheen B, Downs E, Roy RJ, Ailawadi G, Upchurch GR, Jr. A novel chronic advanced stage abdominal aortic aneurysm murine model. J Vasc Surg. 2017;66:232–242 e234. doi: 10.1016/j.jvs.2016.07.105

38. Berman AG, Romary DJ, Kerr KE, Gorazd NE, Wigand MM, Patnaik SS, Finol EA, Cox AD, Goergen CJ. Experimental aortic aneurysm severity and growth depend on topical elastase concentration and lysyl oxidase inhibition. Sci Rep. 2022;12:99. doi: 10.1038/s41598-021-04089-8

39. Barisione C, Charnigo R, Howatt DA, Moorleghen JJ, Rateri DL, Daugherty A. Rapid dilation of the abdominal aorta during infusion of angiotensin II detected by noninvasive high-frequency ultrasonography. J Vasc Surg. 2006;44:372–376. doi: S0741-5214(06)00785-3

40. Rateri DL, Howatt DA, Moorleghen JJ, Charnigo R, Cassis LA, Daugherty A. Prolonged infusion of angiotensin II in apoE(-/-) mice promotes macrophage recruitment with continued expansion of abdominal aortic aneurysm. Am J Pathol. 2011;179:1542–1548. doi: S0002-9440(11)00548-7

41. Al-Tamimi M, Mu FT, Moroi M, Gardiner EE, Berndt MC, Andrews RK. Measuring soluble platelet glycoprotein VI in human plasma by ELISA. Platelets. 2009;20:143–149. doi: 10.1080/09537100802710286

42. Benson TW, Conrad KA, Li XS, Wang Z, Helsley RN, Schugar RC, Coughlin TM, Wadding-Lee C, Fleifil S, Russell HM, et al. Gut Microbiota-Derived Trimethylamine N-Oxide Contributes to Abdominal Aortic Aneurysm Through Inflammatory and Apoptotic Mechanisms. Circulation. 2023;147:1079–1096. doi: 10.1161/circulationaha.122.060573

43. Schmidt RA, Morrell CN, Ling FS, Simlote P, Fernandez G, Rich DQ, Adler D, Gervase J, Cameron SJ. The platelet phenotype in patients with ST-segment elevation myocardial infarction is different from non-ST-segment elevation myocardial infarction. Transl Res. 2018;195:1–12. doi: 10.1016/j.trsl.2017.11.006

44. Cameron SJ, Ture SK, Mickelsen D, Chakrabarti E, Modjeski KL, McNitt S, Seaberry M, Field DJ, Le NT, Abe J, et al. Platelet Extracellular Regulated Protein Kinase 5 Is a Redox Switch and Triggers Maladaptive Platelet Responses and Myocardial Infarct Expansion. Circulation. 2015;132:47–58. doi: 10.1161/CIRCULATIONAHA.115.015656

45. Barkalow FJ, Barkalow KL, Mayadas TN. Dimerization of P-selectin in platelets and endothelial cells. Blood. 2000;96:3070–3077.

46. Offermanns S. Activation of platelet function through G protein-coupled receptors. Circ Res. 2006;99:1293–1304. doi: 10.1161/01.Res.0000251742.71301.16

47. Isselbacher EM, Preventza O, Hamilton Black J, 3rd, Augoustides JG, Beck AW, Bolen MA, Braverman AC, Bray BE, Brown-Zimmerman MM, Chen EP, et al. 2022 ACC/AHA Guideline for the Diagnosis and Management of Aortic Disease: A Report of the American Heart Association/American College of Cardiology Joint Committee on Clinical Practice Guidelines. Circulation. 2022;146:e334–e482. doi: 10.1161/cir.0000000000001106

48. Zhu C, Leach JR, Tian B, Cao L, Wen Z, Wang Y, Liu X, Liu Q, Lu J, Saloner D, et al. Evaluation of the distribution and progression of intraluminal thrombus in abdominal aortic aneurysms using high-resolution MRI. J Magn Reson Imaging. 2019;50:994–1001. doi: 10.1002/jmri.26676

49. Olson JD. D-dimer: An Overview of Hemostasis and Fibrinolysis, Assays, and Clinical Applications. Adv Clin Chem. 2015;69:1–46. doi: 10.1016/bs.acc.2014.12.001

50. Lo RC, Schermerhorn ML. Abdominal aortic aneurysms in women. J Vasc Surg. 2016;63:839–844. doi: 10.1016/j.jvs.2015.10.087

51. Talvitie M, Stenman M, Roy J, Leander K, Hultgren R. Sex Differences in Rupture Risk and Mortality in Untreated Patients With Intact Abdominal Aortic Aneurysms. J Am Heart Assoc. 2021;10:e019592. doi: 10.1161/jaha.120.019592

52. Elbadawi A, Omer M, Ogunbayo G, Owens P, 3rd, Mix D, Lyden SP, Cameron SJ. Antiplatelet Medications Protect Against Aortic Dissection and Rupture in Patients With Abdominal Aortic Aneurysms. J Am Coll Cardiol. 2020;75:1609–1610. doi: 10.1016/j.jacc.2020.02.012

53. Morrell CN, Mix D, Aggarwal A, Bhandari R, Godwin M, Owens P, 3rd, Lyden SP, Doyle A, Krauel K, Rondina MT, et al. Platelet olfactory receptor activation limits platelet reactivity and growth of aortic aneurysms. J Clin Invest. 2022;132. doi: 10.1172/jci152373

54. Touat Z, Ollivier V, Dai J, Huisse MG, Bezeaud A, Sebbag U, Palombi T, Rossignol P, Meilhac O, Guillin MC, et al. Renewal of mural thrombus releases plasma markers and is involved in aortic abdominal aneurysm evolution. Am J Pathol. 2006;168:1022–1030. doi: 10.2353/ajpath.2006.050868

55. Dai J, Louedec L, Philippe M, Michel JB, Houard X. Effect of blocking platelet activation with AZD6140 on development of abdominal aortic aneurysm in a rat aneurysmal model. J Vasc Surg. 2009;49:719–727. doi: 10.1016/j.jvs.2008.09.057

56. Nieswandt B, Bergmeier W, Schulte V, Rackebrandt K, Gessner JE, Zirngibl H. Expression and function of the mouse collagen receptor glycoprotein VI is strictly dependent on its association with the FcRgamma chain. J Biol Chem. 2000;275:23998–24002. doi: 10.1074/jbc.M003803200

57. Lockyer S, Okuyama K, Begum S, Le S, Sun B, Watanabe T, Matsumoto Y, Yoshitake M, Kambayashi J, Tandon NN. GPVI-deficient mice lack collagen responses and are protected against experimentally induced pulmonary thromboembolism. Thromb Res. 2006;118:371–380. doi: 10.1016/j.thromres.2005.08.001

58. Mangin P, Yap CL, Nonne C, Sturgeon SA, Goncalves I, Yuan Y, Schoenwaelder SM, Wright CE, Lanza F, Jackson SP. Thrombin overcomes the thrombosis defect associated with platelet GPVI/FcRgamma deficiency. Blood. 2006;107:4346–4353. doi: 10.1182/blood-2005-10-4244

59. Schulte V, Reusch HP, Pozgajová M, Varga-Szabó D, Gachet C, Nieswandt B. Two-phase antithrombotic protection after anti-glycoprotein VI treatment in mice. Arterioscler Thromb Vasc Biol. 2006;26:1640–1647. doi: 10.1161/01.ATV.0000225697.98093.ed

60. Savage B, Saldívar E, Ruggeri ZM. Initiation of platelet adhesion by arrest onto fibrinogen or translocation on von Willebrand factor. Cell. 1996;84:289–297. doi: 10.1016/s0092-8674(00)80983-6

61. Moog S, Mangin P, Lenain N, Strassel C, Ravanat C, Schuhler S, Freund M, Santer M, Kahn M, Nieswandt B, et al. Platelet glycoprotein V binds to collagen and participates in platelet adhesion and aggregation. Blood. 2001;98:1038–1046. doi: 10.1182/blood.v98.4.1038

62. Sarratt KL, Chen H, Zutter MM, Santoro SA, Hammer DA, Kahn ML. GPVI and alpha2beta1 play independent critical roles during platelet adhesion and aggregate formation to collagen under flow. Blood. 2005;106:1268–1277. doi: 10.1182/blood-2004-11-4434

63. Cosemans JM, Kuijpers MJ, Lecut C, Loubele ST, Heeneman S, Jandrot-Perrus M, Heemskerk JW. Contribution of platelet glycoprotein VI to the thrombogenic effect of collagens in fibrous atherosclerotic lesions. Atherosclerosis. 2005;181:19–27. doi: 10.1016/j.atherosclerosis.2004.12.037

64. Penz S, Reininger AJ, Brandl R, Goyal P, Rabie T, Bernlochner I, Rother E, Goetz C, Engelmann B, Smethurst PA, et al. Human atheromatous plaques stimulate thrombus formation by activating platelet glycoprotein VI. Faseb j. 2005;19:898–909. doi: 10.1096/fj.04-2748com

65. Alshehri OM, Hughes CE, Montague S, Watson SK, Frampton J, Bender M, Watson SP. Fibrin activates GPVI in human and mouse platelets. Blood. 2015;126:1601–1608. doi: 10.1182/blood-2015-04-641654

66. Goebel S, Li Z, Vogelmann J, Holthoff HP, Degen H, Hermann DM, Gawaz M, Ungerer M, Münch G. The GPVI-Fc fusion protein Revacept improves cerebral infarct volume and functional outcome in stroke. PLoS One. 2013;8:e66960. doi: 10.1371/journal.pone.0066960

67. Ungerer M, Li Z, Baumgartner C, Goebel S, Vogelmann J, Holthoff HP, Gawaz M, Münch G. The GPVI-Fc fusion protein Revacept reduces thrombus formation and improves vascular dysfunction in atherosclerosis without any impact on bleeding times. PLoS One. 2013;8:e71193. doi: 10.1371/journal.pone.0071193

68. Uphaus T, Richards T, Weimar C, Neugebauer H, Poli S, Weissenborn K, Imray C, Michalski D, Rashid H, Loftus I, et al. Revacept, an Inhibitor of Platelet Adhesion in Symptomatic Carotid Stenosis: A Multicenter Randomized Phase II Trial. Stroke. 2022;53:2718–2729. doi: 10.1161/strokeaha.121.037006

69. Mayer K, Hein-Rothweiler R, Schüpke S, Janisch M, Bernlochner I, Ndrepepa G, Sibbing D, Gori T, Borst O, Holdenrieder S, et al. Efficacy and Safety of Revacept, a Novel Lesion-Directed Competitive Antagonist to Platelet Glycoprotein VI, in Patients Undergoing Elective Percutaneous Coronary Intervention for Stable Ischemic Heart Disease: The Randomized, Double-blind, Placebo-Controlled ISAR-PLASTER Phase 2 Trial. JAMA Cardiol. 2021;6:753–761. doi: 10.1001/jamacardio.2021.0475

70. Jones B, Tonniges JR, Debski A, Albert B, Yeung DA, Gadde N, Mahajan A, Sharma N, Calomeni EP, Go MR, et al. Collagen fibril abnormalities in human and mice abdominal aortic aneurysm. Acta Biomater. 2020;110:129–140. doi: 10.1016/j.actbio.2020.04.022

71. Wichaiyo S, Parichatikanond W, Rattanavipanon W. Glenzocimab: A GPVI (Glycoprotein VI)-Targeted Potential Antiplatelet Agent for the Treatment of Acute Ischemic Stroke. Stroke. 2022;53:3506–3513. doi: 10.1161/strokeaha.122.039790

72. Daye D, Walker TG. Complications of endovascular aneurysm repair of the thoracic and abdominal aorta: evaluation and management. Cardiovasc Diagn Ther. 2018;8:S138–s156. doi: 10.21037/cdt.2017.09.17

