## Supplemental Methods and Figures for "Glycoprotein VI is Critical for the Detection and Progression of Abdominal Aortic Aneurysms"

Department of Internal Medicine
[1]Division of Cardiovascular Health & Disease
[5]Pathobiology and Molecular Medicine Graduate Program
University of Cincinnati College of Medicine
Cincinnati, OH 45267-0542, USA

Department of Medicine
[2]Division of Epidemiology, Vanderbilt Genetics Institute, Institute of Medicine and Public Health
[3]Vanderbilt-Ingram Cancer Center
[4]Division of Nephrology and Hypertension, Center for Kidney Disease
Vanderbilt University Medical Center
Nashville, TN 37203

[6]Division of Genome Science and Cancer
John Curtin School of Medical Research
The Australian National University
Canberra, ACT, Australia

[7]Heart and Vascular Institute, Department of Cardiovascular Medicine, Section of Vascular Medicine
[8] Lerner Research Institute, Department of Cardiovascular and Metabolic Sciences, [9] Taussig Cancer Institute, Department of Hematology
Cleveland Clinic Foundation
Cleveland, OH 44195, USA

[10]Department of Surgical Sciences, Vascular Surgery
Uppsala University
Uppsala, Sweden

[11]Department of Surgical and perioperative Sciences, Surgery
Umeå, University
Umeå, Sweden

[12]Department of Surgery, Division of Vascular Surgery
University of Rochester School of Medicine
Rochester, NY 14623, USA

[13]Division of Hematology and Oncology
Department of Medicine
UNC Blood Research Center
University of North Carolina at Chapel Hill
Chapel Hill, NC 27599, USA

*Shared first authorship; equal contributions

**Condensed Title:** GPVI is critical to AAA progression

**Address for Correspondence:**

A. Phillip Owens III, PhD

University of Cincinnati

231 Albert Sabin Way ML: 0542

Cincinnati, OH 45267-0542

**COMPLETE MATERIALS AND METHODS:**

*Mouse Study approvals*

Most mouse studies were performed under protocol #15-01-29-01 and continued with protocol #20-11-05-02 with approval and in accordance within the guidelines of the University of Cincinnati Institutional Animal Care and Use. Select mouse studies were performed at the University of North Carolina at Chapel Hill, with approval from the University of North Carolina at Chapel Hill Institutional Animal Care and Use Committee.

*Mice and diet*

Male low-density lipoprotein receptor deficient (*Ldlr*$^{-/-}$; B6.129S7-*Ldlr*$^{tm1Her}$/J; stock: 2207) and *C57BL/6J* (stock: 000664) mice utilized in our studies were obtained from The Jackson Laboratory (Bar Harbor, MA). To induce hypercholesterolemia, *Ldlr*$^{-/-}$ mice were fed a diet enriched with saturated milk fat (21% wt/wt) and cholesterol (0.15% wt/wt, Harlan Teklad diet TD.88137 produced by Purina) for 1 week prior to AngII infusion and throughout the duration of this infusion. C57BL/6J mice were fed a standard chow diet (LabDiet PicoLab® Rodent Diet 20 5053) throughout the course of experimentation. Food and water were given ad libitum. Mice were housed in cages of 2 – 4 with the AngII model, while mice were singly housed in elastase experiments to minimize surgical complications.

*Angiotensin II model of AAA*

Male Ldlr-/- (n = ??) mice underwent infusion of sterile saline or AngII (1,000 ng/kg/min) via a subcutaneously placed osmotic minipump (Alzet Corp) for 28 days, as previously described.In brief, mice were anesthetized with 3.0% isoflurane and prophylactically treated with subcutaneous injection of Buprenorphine SR at a dose of 1mg/kg as analgesic. An Azlet osmotic mini-pump (Model 2004) was implanted subcutaneously into the interscapular space. The pump released either sterile PBS or AngII (Bachem, Product No. 4006473) at a rate of 1,000 ng/kg/min for up to 28 days as previously described.[34] After the infusion period, mice were anesthetized and euthanized by collection of blood and organs.

*Topical elastase model of AAA*

Male *C57BL/6J* mice (n=20) between 8 – 12 weeks of age were anesthetized with 2.5% isoflurane and prophylactically treated with subcutaneous injection of Buprenorphine SR at a dose of 1mg/kg. A ventral midline laparotomy was performed on the mice, exposing the infrarenal aorta. A 4mm piece of sterile Whatman filter paper was placed on the adventitia of the aorta and a sterile solution of 10mg/mL elastase solution (Millipore Sigma, E7885-5, Lot #: SLCL3617) is dropped onto the filter paper and incubated for 5 minutes.[35,36]

The filter paper was then removed, abdomen washed with sterile saline, intestines returned to normal position, and the incision closed in two layers with the body wall closed with absorbable suture and skin with non-absorbable sutures. After 28 days, mice were anesthetized and euthanized by collection of blood and organs.

*Measurements of abdominal aortic diameters*

Abdominal aortas were visualized *in vivo* with high-frequency ultrasound (Vevo 2100, VisualSonics, Toronto, ON, Canada) on day 0, 7, 14, 21 and 28 as described previously.[37,38] Luminal diameters were measured on images with the maximal dilation. Represented diameters in publication are from in vivo ultrasound measurements. For ex vivo measurement, aortas were dissected and cleaned before imaging (Nikon SMZ800N dissecting microscope with 16MP camera) in range of a ruler with standard millimeter measurements. The maximal outer width of the aorta was determined with Nikon Imaging Software (NIS Elements Documentation Version 4.4). AAA was defined as a minimal outer diameter ≥ 1.2 mm. Quantification was verified by a second observer blinded to study groups.[34]

*ELISAs and cytokine array*

A commercially available kit was used to measure D-dimer (Asserachrom; Diagnostica Stago) following manufacturer specifications. Human sGPVI was measured using a solid-phase rabbit anti-GPVI polyclonal antibody and fluid-phase murine anti-GPVI monoclonal antibody (1A12) as previously described.[39]

*JAQ1 antibody intervention*

Mice in both the AngII (*Ldlr*$^{-/-}$ *mice*) and elastase (*C57BL/6J mice*) model were screened for the development of a AAA (50% increase in aortic diameter from baseline) via ultrasound 14 days post model initiation. Mice were then blindly randomized to either the JAQ1 intervention or control groups normalized for aortic diameter. Once randomized, each mouse received a one-time 250 µl intraperitoneal injection of 50µg of JAQ1 GPVI neutralizing antibody (Emfret Analytics, Germany, Catalog: M011-0) or 50µg rat IgG2a placebo as described previously.[29]

*Mouse platelet isolation and analysis by flow cytometry*

Approximately 1 mL of blood was drawn from the IVC into syringe containing 100 µL of 3.2% sodium citrate. Blood was diluted with 2 volumes of Tyrode's buffer then centrifuged at 200g for 4 minutes. Platelet rich plasma (PRP) layer was gently pipetted off. PRP was treated with acid citrate dextrose, prostaglandin I2 (Fisherscientific, AAJ67115MCR) and apyrase (Sigma-Aldrich, A6535) and allowed to rest for 30 minutes. PRP was then centrifuged at 2,000g for 4 minutes, forming a platelet pellet. After resuspension in 100 µL of Tyrode's buffer, platelets were exposed to several concentrations of platelet agonist thrombin (Cayman Chemical, 13188) or convulxin (Cayman Chemical, 19082) in the presence of platelet specific marker CD41 antibody (BioLegend, 133934) and activation marker CD62 P-selectin antibody (BD Biosciences, 553744). Platelets were then fixed in 2% formalin for 10 minutes. Platelet isolation was confirmed by staining Alexa Fluor 647 anti-mouse CD41 antibody (BioLegend, 133934). CD41 positive cells were analyzed for platelet surface activation marker FITC rat anti-mouse CD62P (BD Biosciences, 553744) for P-selectin using CytoFLEX flow cytometer (Beckman Coulter) at 100K events per sample. P-selectin mean fluorescence was quantified using the geometric mean.

*Mouse Platelet Counts*

A 15 µL aliquot of whole blood was used to assess platelet count via a veterinary hematology analyzer (Heska Element HT5). Platelet count is reported as PLT x $10^3$/µL.

*Histological processing of abdominal aortas*

Histological processing and staining of mouse aortas was completed as previously described.[40] Briefly, a section of abdominal aorta was embedded in optimal cutting temperature compound and frozen at -20°C. Tissue was sectioned on a cryostat at 10 µm and underwent picrosirius red or CD68 staining to quantify Type I Collagen or macrophages, respectively. Picrosirius red (PolyScientific Cat No 24901-500) staining was performed following manufacturer instructions and imaged using a 4X objective using bright and polarized light. Type I collagen was measured by comparing red intensity to total intensity in ImageJ software. To stain for CD68, sections were fixed in ice cold acetone for 20 minutes and permeabilized in 0.06% solution of triton-X 100 (Sigma-Aldrich Cat No T8787) in TBS for 30 minutes at room temperature followed by three washes in 1X TBS. Sections were blocked for 1.5 hrs with 4% Normal Goat Serum (Life Technologies REF 500622) in TBS. Primary antibody rat anti-mouse CD68 Alexa Fluor® (BioRad Cat No MCA1957A647) was diluted 1:500 in blocking solution and incubated at 4°C overnight. The next day, samples were incubated in secondary antibody Alexa Fluor 488 (Thermo Scientific Cat No A-11006) diluted 1:500 in blocking solution for 1 hour at room temperature. Invitrogen ProLong™ Gold antifade reagent with DAPI (Thermo Scientific REF P36935) was used to mount the sections. Stellaris 8 Confocal microscope captured images using 4X and 10X objectives and CD68+ percent area was calculated in ImageJ comparing fluorescent area to total area.

*Human aneurysm samples*

Human AAA tissue was obtained from four patients (4 males) aged 65 ± 7.7 years (mean ± SD; range 45 to 71 years) undergoing open aneurysm repair (between 2015 and 2018) at the University of Rochester Medical Center in Rochester, NY under an approved University of Rochester Cardiovascular Tissue Bank Internal Review Board Application (RSRB 00036669). The patient criteria for open surgical repair were defined as AAA diameter exceeding 55 mm for males, AAA diameters between 50 and 55 mm for females, rapidly expanding aortic diameters (≥ 5 mm in 6 months with a minimum diameter of 40 mm), or symptoms attributable to AAA and AAA rupture. Mural thrombi (easily separated from the aortic wall) were collected during surgery, along with residual aortic wall fragments, and were immediately flash frozen until protein processing. Control tissue was procured from the infrarenal aortas of 5 males aged 58 ± 6.6 years (mean ± SD; range 42 to 65 years) cause of death deemed non-aortic causes. Tissues were collected within 24 hours of death and flash frozen for analysis.

This study complies with the Declaration of Helsinki and was approved by the University of Rochester for the analysis of platelets from healthy individuals, and from patients with AAA as well as blood biomarkers and aortic tissue from recently deceased individuals from non-vascular causes as well as from patients undergoing open aortic reconstruction. The Cleveland Clinic Institutional Review Board approval was gained for the isolation of platelets from patients with AAA.

*RNA processing*

RNA was isolated from human AAA wall, accompanying thrombi, and control aortic walls (described above) tissue with the commercially available NucleoSpin® RNA kit (Macherey-Nagel REF 740955) with several modifications. Tissue was thawed on ice and transferred to BeadBug™ prefilled tubes (Benchmark Scientific Triple-Pure High Impact Zirco nium Beads Item D1032-15). Tubes were filled with 1 mL QIAzol Lysis Reagent (QIAGEN Cat No. 79306) and homogenized with a BeadBug™ Microtube Homogenizer (Benchmark Scientific) after which 200 µL of chloroform were added. Samples were vortexed and left to sit at room temperature for 2 – 3 minutes. Samples were then centrifuged at 12,000 X g for 15 minutes at 4°C. The aqueous phase was removed to new tubes and 150 µL isopropyl alcohol was added to each. Following a 10-minute incubation at room temperature, each lysate was loaded onto a NucleoSpin® RNA Column and eluted to RNA. RNA was quantified with a NanoDrop™ 2000 Spectrophotometer (Thermo Scientific) and samples were stored at -80°C.

*RNA sequencing*

RNAseq was performed by the Cincinnati Children's Gene Expression Core Facility. Next generation sequencing was conducted with an RNA polyA stranded library against homo sapien. Parameters were set at paired ends with 100 bases and were read at a depth of 40 million reads per sample. Data was analyzed using the bioinformatics software CLC Genomics Workbench version 20.0.3 and Ingenuity Pathway Analysis (Qiagen, Hilden, Germany).

*Platelet isolation and immunoblotting*

Healthy patients (n = 4) were volunteers without any known medical history, aneurysmal disease, or on antiplatelet agents were recruited in studies approved by the Institutional Review Board at the Cleveland Clinic. All AAA patient samples (n = 5) were collected by venipuncture from pre-operative patients under serial imaging surveillance. For each subject, blood was drawn by a medical professional into citrate plasma tubes, then centrifuged in a tabletop centrifuge at 1100 rpm for 15 mins. Platelet rich plasma (PRP) well above the buffy coat was decanted and the final platelet centrifugation step at 2600 rpm for 5 mins was conducted with a final concentration of 10 μM $PGI_2$ (in Tris Buffer, pH 9.0). The final washed platelet pellet from one human plasma citrate tube was resuspended in 1000 μL of fresh Tyrode's solution which was diluted 1:10 in fresh Tyrode's solution. This was aliquoted into 100 μL platelet samples before reducing SDS-PAGE and Immunoblotting as described by us previously.[41,42] Blocking buffer was 3% bovine serum albumin/Tris-buffered saline–Tween 20 for 60 minutes at room temperature with agitation. TBST-T Primary anti-GPVI antibody (ABCAM, # ab129019) was used in a 1:4000 dilution in 3% bovine serum albumin/Tris-buffered saline–Tween 20 for 12 hours at 4C with agitation. GAPDH antibody (Cell Signaling Technology #5174S) was used in a 1:000 dilution in 3% bovine serum albumin/Tris-buffered saline–Tween 20 for 12 hours at 4 degrees C with agitation. Secondary antibody (GE Healthcare, Buckinghamshire, UK) was used in a 1:2000 titer in 5% milk/Tris-buffered saline–Tween 20 for 1 hour at room temperature with agitation. Final autoradiographic films (Bioblot BXR, Laboratory Product Sales, Rochester NY) were quantified by densitometry using ImageJ software (National Institutes of Health).

*Human platelet surface GPVI quantification by flow cytometry*

Isolated 100 μL platelet samples were incubated with 1 μL of labeled GPVI antibody (BD Biosciences, Clone HY101, #565241 in the dark for 30 minutes). This reaction was then stopped by adding 100 μL of 2%

formalin to each reaction, and then quantification of platelet surface GPVI was made possible using an Accuri C6 Plus Flow Cytometer (BD Biosciences) at 10K events per sample. Data was then processed through FloJo (Ashland, Oregon). Platelet surface GPVI was quantified using the geometric mean.

*Human Platelet activation of control and AAA patients in response to convulxin*

Blood was drawn by a medical professional into citrate plasma tubes, then centrifuged in a tabletop centrifuge at 1,100 rpm for 15 minutes. Platelet rich plasma (PRP) well above the buffy coat was decanted. Final platelet centrifugation at 2,600 rpm for 5 mins was conducted with a final concentration of 10 µM PGI 2 (in Tris Buffer, pH 9.0). The final washed platelet pellet from one human plasma citrate tube was resuspended in 1000 µL of fresh Tyrode's solution which was diluted 1:10 in fresh Tyrode's solution. Following 15 minutes of agonist stimulation, 1 µL of labeled CD62P (P-selectin) antibody was incubated in the dark for 30 minutes. Samples were fixed in 2% formalin, then platelet surface P-selectin was quantified on an Accuri Flow Cytometer (BD Biosciences).

*Research statistics and data representation*

Unless indicated otherwise, all bar, scatter, and line graphs were created with Sigma Plot v.14.5 (SPSS, Chicago, IL). Data are represented as individual data point and mean ± SEM. Data normality was assessed using a Shapiro Wilk test. For two group comparison where normality can be assumed, a Student's t-test was performed, while non-normal data was analyzed with a Mann-Whitney Rank Sum. Statistical significance between multiple groups was assessed by One Way analysis of variance (ANOVA) on Ranks with a Dunn's post hoc, One Way ANOVA with Holm Sidak Post Hoc, or Two-Way ANOVA with Holm Sidak Post Hoc, when appropriate. Statistical significance among groups performed temporally was assessed by either a One-Way Repeated Measures ANOVA (parametric) or Repeated Measures ANOVA on Ranks (non-parametric), where appropriate. Values of $P < 0.05$ were considered statistically significant.

*European Study Population*

We studied participants from a population-based case-control study of patients with abdominal aortic aneurysm (AAA) and healthy controls from the Uppsala region between 2008 and 2016 described previously.[7,40] At time of enrollment, all blood, plasma, and aneurysm measurements were collected and at final follow-up, diameter measurement was taken. For the present study, AAA patients were included if aortic diameter was greater than 30 mm and if follow-up was more than 6 months. Patients with ruptured AAA, coexisting malignant disease, dialysis dependence, recent thromboocclusive disease, antithrombotic therapy, or sub aneurysmal aortic dilations were excluded. Fast growing AAA was defined as a change >2 mm per year in diameter. Details of this study have previously been published.[7,40] A total of 169 AAA cases and 115 controls met the inclusion criteria for the study. Female cases (n=20) were excluded because no appropriate female controls were available. A total of 75 patients with slow-growing AAAs, 74 patients with fast-growing AAAs, and 115 controls were included in the present study. A total of 4 participants were missing D-dimer values and were excluded from the D-dimer analyses.

Ordinal logistic regression was used to estimate the odds ratios (OR) and 95% confidence intervals (CI) for the association between sGPVI and case status and D-dimer and case status. sGPVI was modeled dichotomously (above/below median sGPVI, 38ng/mL) and continuously (log base-2 transformed). D-dimer was modeled dichotomously (above/below median D-dimer, 500 ng/mL) and continuously (log base-2 transformed). In analyses restricted to AAA cases only, linear regression was used to evaluate the associations between sGPVI, D-dimer, and difference in diameter and growth rate of AAA.

Fully adjusted models included age, smoking years, aspirin use, and prevalent cardiovascular disease (CVD), coronary heart disease (CHD), diabetes, hypertension, and chronic obstructive pulmonary disease (COPD). Chained multiple imputation was performed for missing values of height (n=64), weight (n=68), number of cigarettes (n=60), smoking years (n=42), CHD (n=5), CVD (n=2), hypertension (n=3), COPD (n=4), claudication (n=4), renal insufficiency (n=2), statin use (n=3), aspirin use (n=3), and diabetes (n=2). Covariates were chosen *a priori* based on biological plausibility (age, smoking years, hypertension, and COPD) and by forward selection that required a significance level of p-value $\leq$ 0.10 (age, smoking years, aspirin use, CVD, CHD, and diabetes).

To predict the occurrence of fast-growing vs. slow-growing AAA, comparative ROC analysis was performed using sGPVI or D-dimer concentrations from AAA patients only, with slow-growing AAA values being used as the reference condition. ROC curves generated using Sigma Plot v.14.5.

The combined effect of sGPVI and D-dimer on case status was examined using ordinal logistic regression. The unadjusted model included continuous (log base-2 transformed) sGPVI and continuous (log base-2 transformed) D-dimer. The adjusted model included D-dimer, sGPVI, age, smoking, CHD, CVD, diabetes, aspirin use, hypertension, and COPD. The combined effect was calculated using the linear combination of D-dimer and sGPVI. In analyses restricted to AAA cases only, linear regression was used to evaluate the combined effect of sGPVI and D-dimer on difference in diameter and growth rate of AAA.

*American Study Population*

We additionally examined the relationship between sGPVI, D-dimer, and AAA in participants from a case-control study of patients with AAA and healthy controls as a part of a prospective cohort in the United States. Details for this cohort have already been published.[40] Plasma samples and corresponding clinical data were collected at tertiary care centers. Sequential participants were enrolled in the study which consists of sequential stable subjects who underwent elective diagnostic coronary angiography (cardiac catheterization or coronary computed tomography) for evaluation of CAD. Abdominal Aortic Aneurysm (AAA) was defined as a baseline abdominal aortic diameter of 3.0 cm or greater and fast-growing AAA was defined as a change >4 mm per year in diameter. A total of 118 AAA cases (89 slow-growing and 29 fast-growing) and 100 controls were included in the study. Height, weight, smoking, comorbidities, medication use, and aorta diameter were only available in AAA cases.

sGPVI was modeled dichotomously (above/below median sGPVI, 21 ng/mL) and continuously (log base-2 transformed). D-dimer was modeled dichotomously (above/below median D-dimer, 455 ng/mL) and continuously (log base-2 transformed). Ordinal logistic regression was used to estimate the ORs and 95% CIs for the association between sGPVI and case status and D-dimer and case status. In analyses restricted to AAA cases only, linear regression was used to evaluate the associations between sGPVI, D-dimer, and growth rate of AAA. Fully adjusted models included age, race, and sex.

To predict the occurrence of fast-growing vs. slow-growing AAA, comparative ROC analysis was performed using sGPVI or D-dimer concentrations from AAA patients only, with slow-growing AAA values being used as the reference condition. ROC curves generated using Sigma Plot v.14.5.

The combined effects of sGPVI and D-dimer on case status and growth rate of AAA were examined using ordinal logistic regression and linear regression. The unadjusted models included continuous (log base-2 transformed) sGPVI and continuous (log base-2 transformed) D-dimer. The adjusted models included D-dimer, sGPVI, age, race, and sex. The combined effects were calculated using the linear combination of D-dimer and sGPVI.

All analyses for both the European and American study populations, except for ROC curves, were completed in Stata version 16.1 (StataCorp, College Station, TX, USA).

**SUPPLEMENTAL FIGURE LEGENDS:**

**Supplemental Figure 1:** *Human AAA thrombus is enriched for platelet transcripts compared to AAA wall and control wall.* (A) Venn-diagram comparing differentially expressed genes of control wall, AAA wall and AAA thrombus. AAA thrombus vs control and AAA thrombus vs AAA wall contained the majority platelet-specific transcripts. (B) Volcano plot of AAA thrombus vs control aortic wall. Platelet-specific transcripts (red dots) were significantly upregulated in AAA thrombus.

**Supplemental Figure 2:** *Mice with established AngII-induced AAAs treated with JAQ1 antibody were protected from AAA rupture and progression.* *Ldlr*$^{-/-}$ mice with established AngII AAAs were treated with either IgG control or JAQ1 antibody. (A) Intervention with JAQ1 had no discernable impact on body weight. (B) None of the mice treated with Jaq1 antibody died during the study period while 3/9 mice in the control group suffered a rupture-induced death. JAQ1 and control aortas were embedded, cryosectioned, and underwent picrosirius red staining for Type I Collagen and CD68 staining for infiltrating monocytes. (C) Representative images for picrosirius staining (left) and CD68 staining (right) were captured with Leica Stellaris 8 Confocal. (D) Aortas from JAQ1 treated mice had significantly higher percentage of Type I Collagen compared to control (n = 5/group. **, P< 0.01 by Student's t-test). (E) JAQ1 treated mice (n = 5) also had a lower percentage of infiltrating macrophages compared to control (n = 4) (P = 0.092 by Welch's t-test). Platelet count at sacrifice (F) did not vary significantly between JAQ1 and control groups.

**Supplemental Figure 3:** *Mice with established elastase induced AAAs treated with Jaq1 antibody were protected from AAA progression.* *Ldlr*$^{-/-}$ mice with established elastase AAAs were treated with either IgG control or JAQ1 antibody. (A) Body Weight and (B) Platelet count was similar between the two groups. (C) Representative images of picrosirius red staining (left) and CD68 staining (right). JAQ1 treated mice maintained significantly more type I collagen compared to control mice (n = 5/group. **, P<0.01 by Student's t-test). JAQ1 treated mice had a lower percentage of infiltrating macrophages compared to control mice (n = 5/group. P = 0.333 by Student's t-test).

#### Supplemental Figure 1

A.

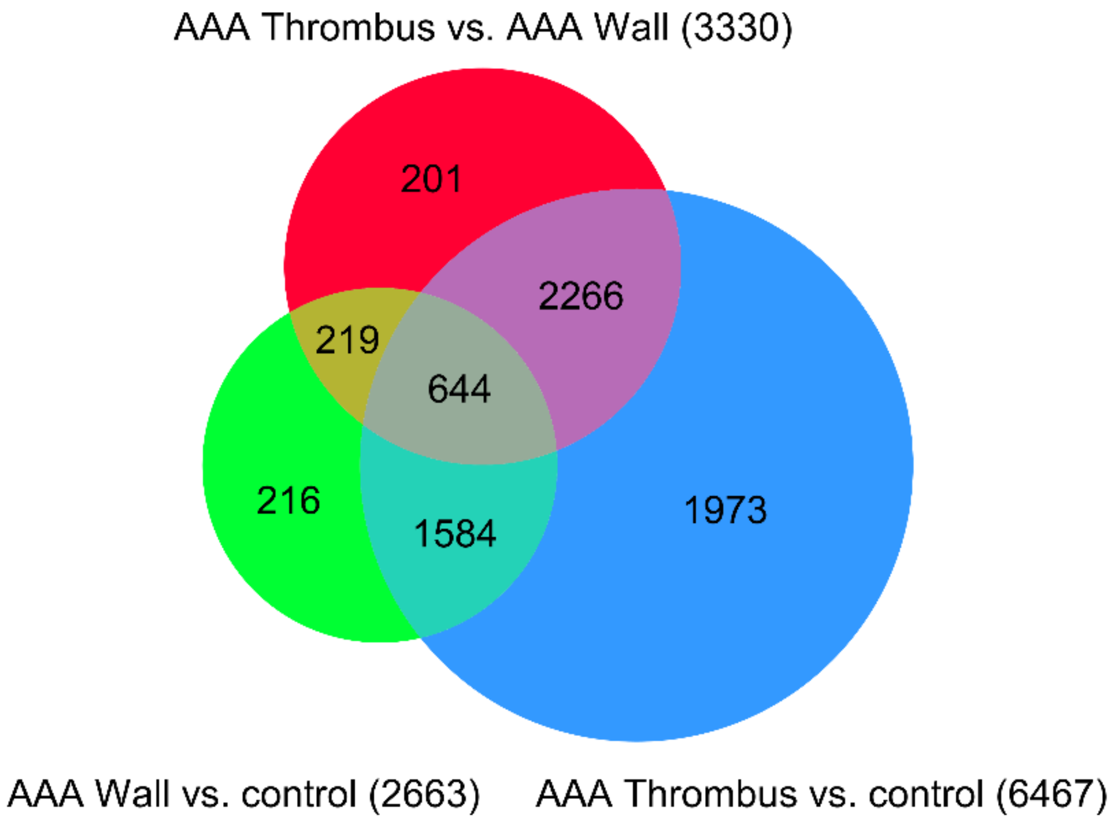


B.

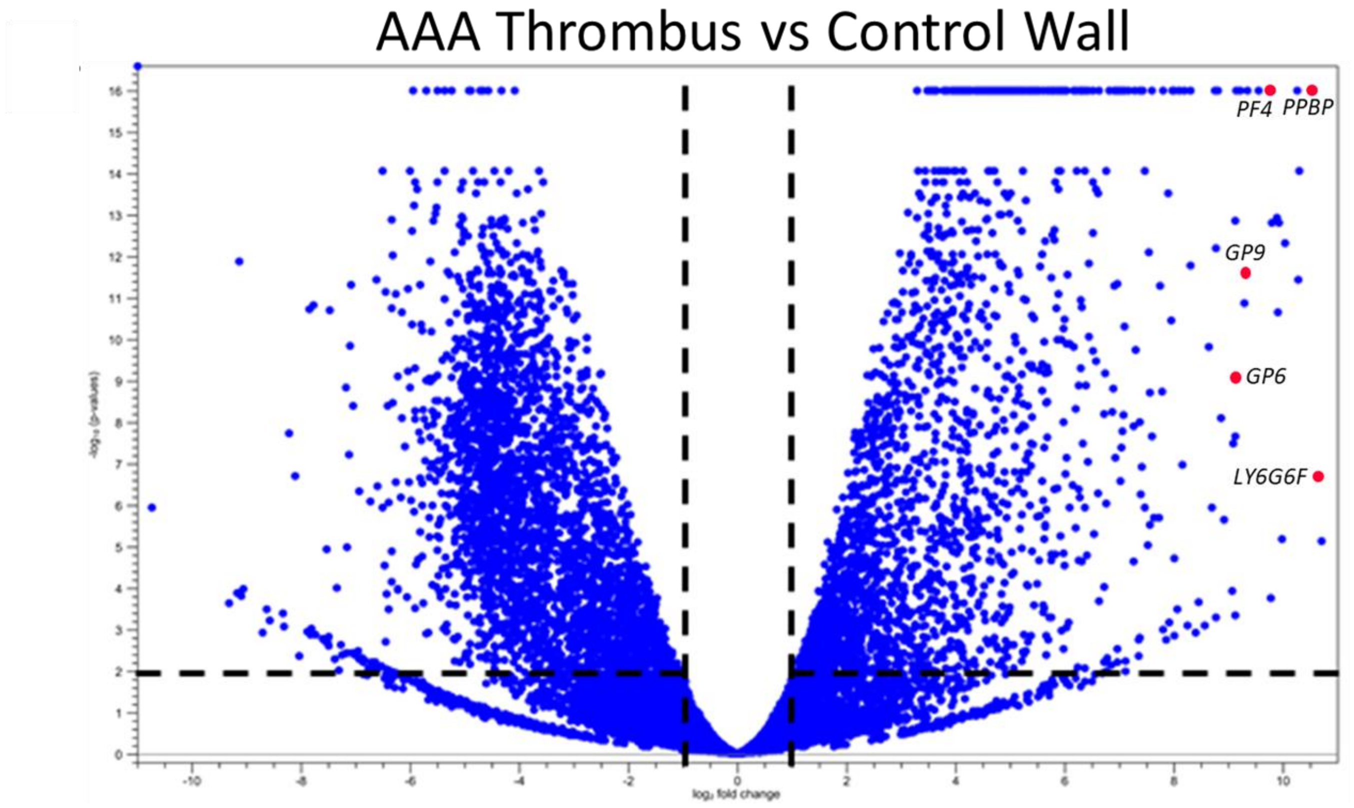

### Supplemental Figure 2

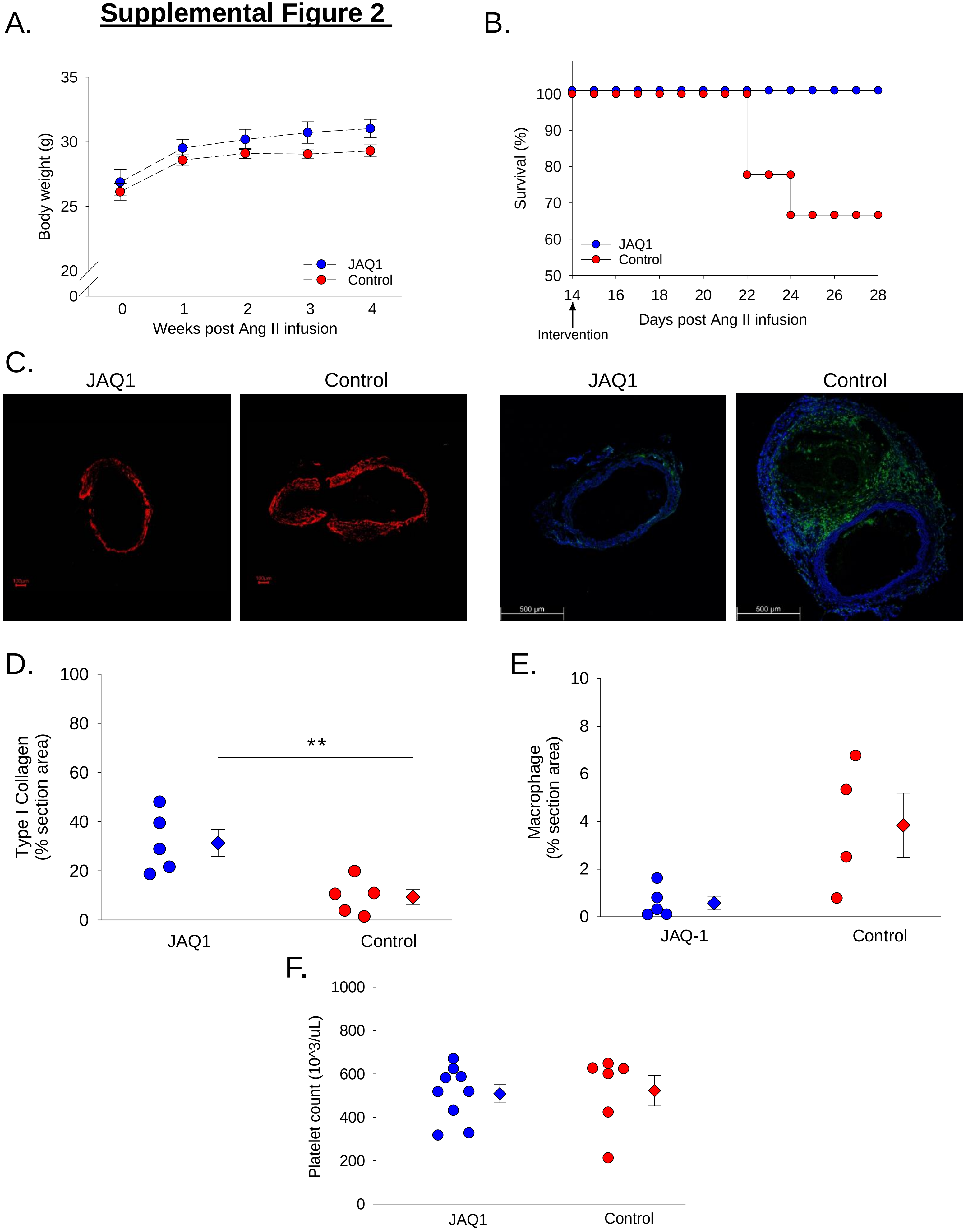

#### Supplemental Figure 3

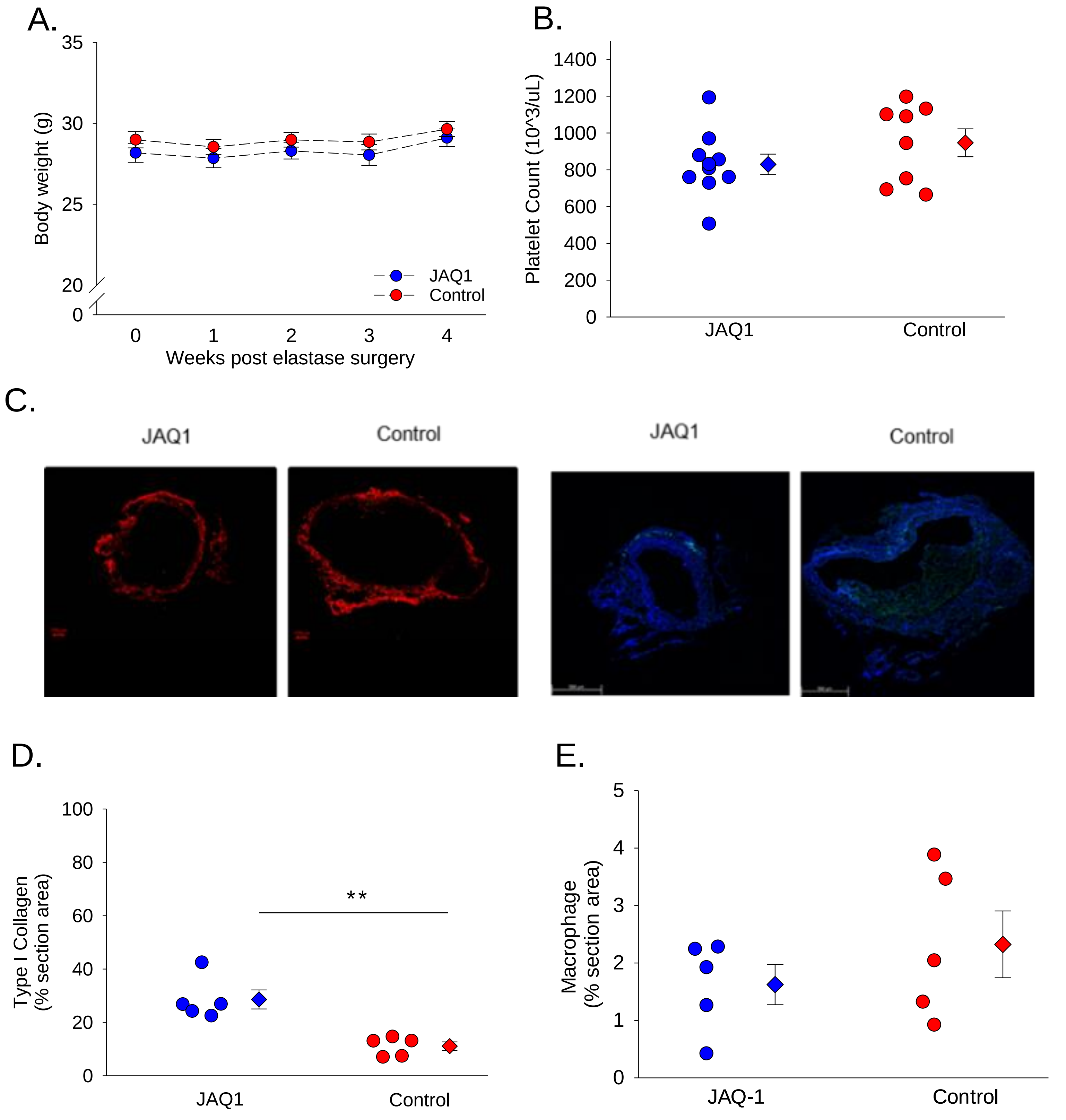
